# *Toxoplasma gondii* infection misdirects placental trophoblast lineage specification

**DOI:** 10.1101/2024.09.10.612241

**Authors:** Leah F. Cabo, Liheng Yang, Mingze Gao, Rafaela J. da Silva, NyJaee N. Washington, Sarah M. Reilly, Christina J. Megli, Carolyn B. Coyne, Jon P. Boyle

## Abstract

Pregnancy is a critical point of vulnerability to infection and other insults that could compromise proper fetal development. The placenta acts as a protective and nutrient-permeable barrier to most infectious agents, but a few are capable of bypassing its defenses. Remarkably little is known about how exposure to these select pathogens might impact ongoing placental development. Here we demonstrate that *Toxoplasma gondii* entirely misdirects the developmental program of trophoblast stem cells. Infection of progenitor cytotrophoblasts prevents fusion and differentiation to infection-resistant syncytiotrophoblast. Rather, *T. gondii* elicits a unique transcriptional identity that polarizes cytotrophoblasts to the infection-permissive extravillous trophoblast fate. Strong evidence of developmental disruption is found in multiple orthogonal models, including trophoblast stem cells, trophoblast organoids, and chorionic villi. Manipulation of cell fate by the parasite is most dramatic in trophoblast organoids, where we see robust outgrowth of HLA-G(+) extravillous trophoblasts. Collectively, these data show that *Toxoplasma* antagonizes differentiation of an infection-resistant cell type by inducing formation of an infection-permissive cell type, therefore potentiating its own transmission to the fetus.

## INTRODUCTION

Congenitally transmitted pathogens account for some of the most common causes of cryptic fetal illness. Pathogens that can cross from mother to fetus during pregnancy are broadly categorized as members of the TORCH (***T****oxoplasma gondii*, **O**ther, **R**ubella, **C**ytomegalovirus, and **H**erpesvirus) group^1^. This includes *Toxoplasma gondii*, a single-celled, intracellular, eukaryotic parasite with globally distributed incidence of congenital transmission^2,3^. Among the potential manifestations of clinical toxoplasmosis, congenital transmission can be particularly severe given fetal vulnerability to parasite-induced pathology^4,5^.

The trophoblastic layers of the human placenta are a primary biological barrier that pathogens must overcome to infect the fetus. The functional unit of the human placenta, the chorionic villi, are comprised of three canonical populations of placental trophoblast. Progenitor cytotrophoblasts (CTB) give rise to differentiated extravillous trophoblasts (EVT) or fused syncytiotrophoblasts (STB), each population with its own role in the physiological function and maintenance of the placenta. CTB are the largest population of trophoblast and pioneer the organ’s establishment and expansion during early to mid-gestation. As the placenta develops, CTB continuously differentiate to supply the terminally differentiated STB and EVT layers.

The syncytiotrophoblast (STB) makes up a large, fused, multinucleated cell layer that lines the surface of chorionic villi. Previous work shows that STB autonomously resist *T. gondii* infection, one of the few known human cell types to do so^6,7,8^. Both primary and *in vitro*-derived STB restrict *T. gondii* at the point of the parasite’s attachment to the cell surface^6,8^. These findings are in line with the known modalities through which STB restrict other congenital pathogens like *Listeria monocytogenes*^9,10^, cytomegalovirus^11^, and herpes simplex virus^12^, suggesting pathogen restriction is one of its primary functions.

Extravillous trophoblasts (EVT) are mesenchymal cells that invade from the distal end of the villous tree core deeply into the maternal decidual tissue. EVT dictate vascular remodeling in the decidua by contributing to the formation and breakdown of plugs in the maternal spiral arteries. They recruit and interact with fetal/maternal macrophages and uterine NK cells, suggesting EVT have local immunomodulatory capacity^13,14^. Several subtypes of EVT have been identified with varied invasiveness^15,16^. The spectrum of functional and transcriptional EVT types characterized likely represent subpopulations that have differing roles in organ establishment, immunological communication and microvasculature remodeling. Studies suggest that the earliest EVT arise from *ITGA2+* CTB at the distal end of the villous tree, a population known as cell column cytotrophoblasts (CCC)^17^. Like CTB, EVT are known to be generally permissive to infection by *Toxoplasma* and other pathogens, therefore this cell population could be a meaningful portal for pathogen entry into the fetal compartment^7,18^.

*Toxoplasma* massively alters the transcriptional landscape of its host cells.^19,20^ Simultaneously, the trophoblast developmental program is tightly regulated to maintain proper proportions of each cell lineage. *Toxoplasma* has a repertoire of protein effectors that have been shown to alter host cell biology specifically to propagate parasite replication and survival^21,22,23^. There is evidence in other systems that transcriptional changes during infection specifically disrupt cell differentiation kinetics, including in *Plasmodium falciparum*^24,25^ and *Trypanosoma cruzi*^26,27^, parasites that, like *Toxoplasma gondii,* are teratogenic intracellular parasites.

In this study, we use a tractable placenta-derived bipotent stem cell system, primary tissue-derived trophoblast organoids, and third trimester chorionic villi to investigate how *Toxoplasma gondii* infection misdirects trophoblast cell fate. Orthogonal analyses reveal that *T. gondii* infection antagonizes STB fusion and development in trophoblast stem cells through induction of epithelial to mesenchymal transition. Activation of this distinct transcriptional program promotes EVT differentiation at the cost of STB development. This infection-induced fate polarization may predicate parasite dissemination to the fetus by reducing infection-resistant STB coverage and creating a pro-invasive environment that supports EVT formation.

## RESULTS

### Hormone secretion and STB marker expression of *Tg-*infected TS-CTB is diminished over time

First-trimester placenta-derived human trophoblast stem cells (henceforth: TS-CTB, TS-STB, and TS-EVT) recapitulate much of the transcriptional and functional identity of their *in vivo* counterparts^6,28,29,30,31^. TS-CTB, TS-STB, and TS-EVT stain appropriately with their cell-identity markers ITGA6, SDC1, and HLA-G respectively (**Figure 1A**). By manipulating culture conditions as previously published, TS cells are used to replicate terminally and intermediately differentiated placental trophoblasts to model their interactions with *Toxoplasma* during congenital infection.

**Figure 1:**
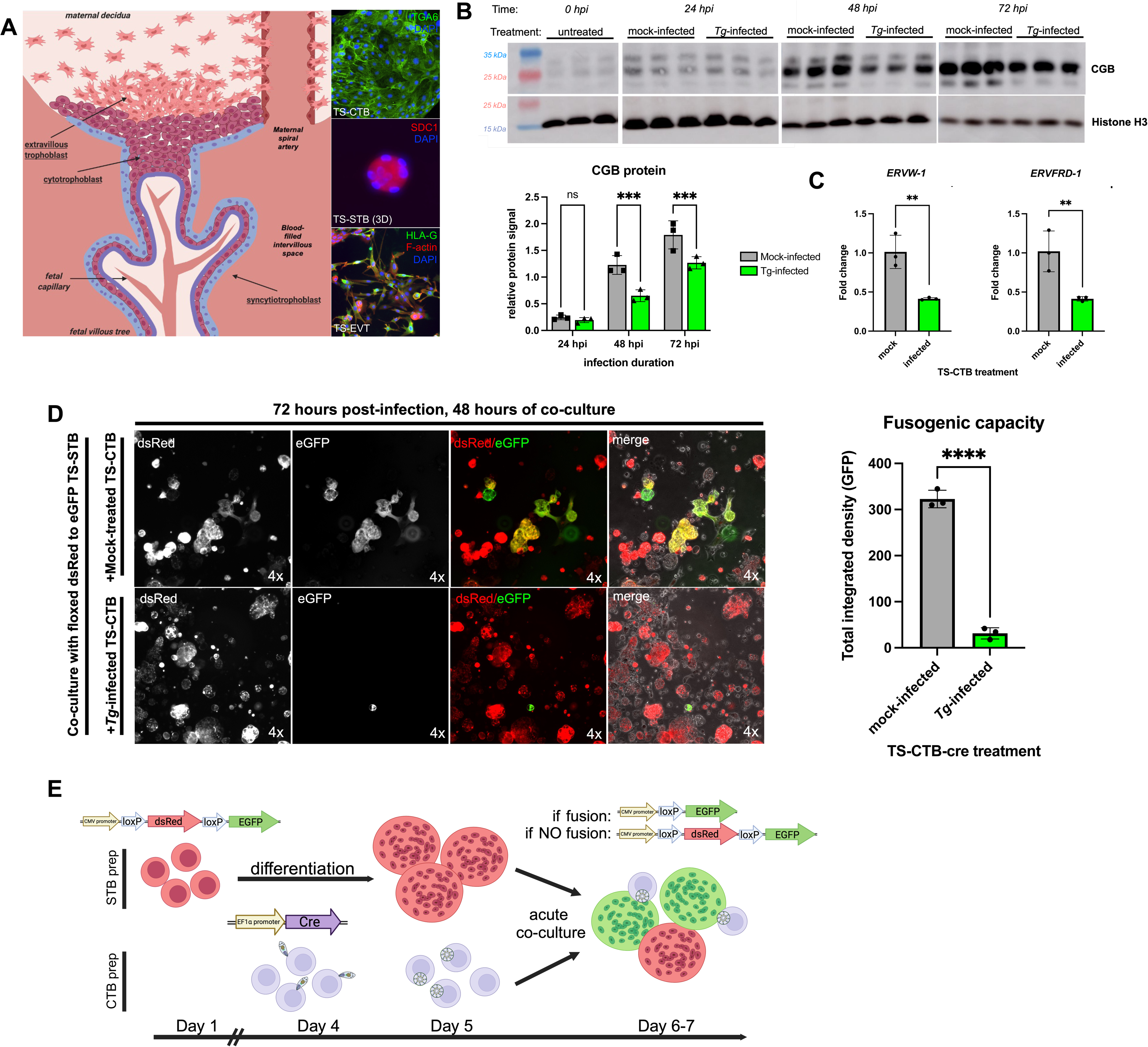
*T. gondii* infection antagonizes STB fate and function. (A) Model of trophoblastic interface between maternal and fetal blood supply. Representative images of TS-CTB, TS-STB (3D) and TS-EVT each stained with cell type-specific markers. (B) Western blot of whole cell lysate from mock-infected and *Tg-*infected TS-CTB at indicated time points. Proteins visualized using hCG polyclonal antibody. Molecular weights of observed bands likely indicate detection of multiple CGB isoforms. Protein quantified using integrated density of observed bands normalized to histone H3 loading control within timepoint. Significance determined with two-way ANOVA with Tukey’s multiple comparisons test. All comparisons were performed but just comparisons of interest are plotted. (C) Syncytin 1 (ERVW-1) and syncytin 2 (ERVFRD-1) transcripts quantified via qPCR of mock-infected versus *Tg-*infected TS-CTB. Housekeeping gene is *GAPDH*. Points represent three averaged technical replicates of one biological replicate. Results are representative of >2 independent experiments. Significance determined by one-sample two-tailed t-test using linear ΔCT values and results for significance projected onto fold change. (D) Representative images of flox dsRed to eGFP TS-STB co-culture with either mock-infected or *Tg-*infected TS-CTB-cre. Red and green channels projected singularly in greyscale or combinatorically in color. Quantification of GFP across conditions by integrated density and normalized to total cell area. Measurements collected at conclusion of 48 hour co-culture. Each point represents an average of 4 fields of view at 4x magnification per single biological replicate for a total of three biological replicates per condition. Results are representative of two independent experiments. (E) Experimental schematic of fusion arrest assay with transgenic cell lines. For all, * p < 0.05, ** p < 0.01, *** p < 0.001, **** p < .0001.

The STB population mediates pathogen resistance in the placenta through both physical and immunological mechanisms^6,7,9,32,33^. Several key biological changes occur during differentiation of CTB to STB, though it has yet to be determined which directly confer pathogen resistance. We hypothesized that if *T. gondii* infection (*Tg-*infection) antagonizes STB formation similarly to other congenital pathogens^34,35^, we would detect defects in two essential indicators of STB formation: hormone secretion and cell fusion.

Of the hormones secreted by STB, pregnancy-maintaining human chorionic gonadotrophin (hCG) is among the most essential. hCG is formed through the dimerization of two protein subunits, CGA and CGB. CGA is among the most abundant transcripts expressed by placental trophoblasts as it is common to all gonadotropic hormones^36,37^. Meanwhile, CGB expression is more restricted to STB and STB precursors, making it a useful indicator of differentiation. To investigate how *Tg-*infection impacts hormone production, we used Western blot to detect CGB expression in whole cell lysates at various time points. At 0 hpi, untreated TS-CTB show appropriately low levels of CGB protein. At 24 hpi, there is no significant difference between CGB of mock- and *Tg-*infected TS-CTB. AT 48 and 72 hpi, we detect a significant difference in CGB detected between mock- and *Tg-*infected TS-CTB (**Figure 1B**). In two separate experimental replicates at a single time point, CGB is almost entirely absent from *Tg-*infected TS-CTB (**Figure S1A**). Collectively, these data indicate a temporally regulated loss of CGB expression in *Tg-*infected TS-CTB. As CGB expression and eventual hCG secretion is a core function of STB in early pregnancy, this phenotype suggests that infection prevents or delays TS-CTB differentiation to TS-STB.

Another critical element of STB formation is cell fusion. Throughout pregnancy, underlying CTB fuse into the existing continuous STB layer. Fusion is canonically mediated by ERVW-1 (syncytin 1) and ERVFRD-1 (syncytin 2), both of which mediate creation of the fusion pore and membrane merging^38,39,40^. Increased *ERVW-1* and *ERVFRD-1* serve as both a transcriptional marker and the direct molecular driver of STB differentiation. We detected significant reductions of both *ERVW-1* and *ERVFRD-1* transcript (both 59% reduction) via qPCR after 48 hours of *Tg-* infection in TS-CTB (**Figure 1C**). The reduction of these essential fusogenic genes further indicates suppression of STB fate, and simultaneously, suggests functional arrest in CTB-to-STB fusion during *Toxoplasma* infection.

### *Toxoplasma* infection causes near complete functional arrest of trophoblast fusion

Our transcriptional and Western blot data led to the hypothesis that *T. gondii* infection may also alter or suppress trophoblast fusion. While we see reduced expression of the two primary fusion markers *ERVW-1* and *ERVFRD-1* (**Figure 1C**), this does not functionally assess whether *T. gondii* infection prevents fusion. Therefore, we developed an assay designed to directly assess the physical fusion of *Tg-*infected CTB to mature STB. Using two transgenic TS-CTB lines, one with a *loxP*-flanked *dsRed* and a downstream *eGFP* construct driven by a CMV promoter and another with cre-recombinase driven by an EF-1 promoter, we conducted acute co-culture experiments with either mock- or *Tg-*infected TS-CTB-cre and floxed dsRed STB (**Figure 1E**). If fusion were to occur between TS-CTB-cre and floxed *dsRed* TS-STB, the cre-recombinase will recombine the *loxP* sites flanking *dsRed*, deleting the *dsRed* gene and moving the CMV promoter proximal to *eGFP* sequence, producing green TS-STB cells that will also gradually lose detectable dsRed (fusion(+)). If fusion does not occur between TS-CTB-cre and floxed *dsRed* TS-STB, the TS-STB will remain exclusively red (fusion(-)) (**Figure 1E**).

When we used this assay and quantified GFP fluorescence integrated density as a proxy for fusion, *Tg-*infected co-cultures showed a near complete loss of fusion capacity compared to mock-infected (**Figure 1D**). Qualitatively, 30-50% TS-STB became fusion(+) and turned green when co-cultured with mock-infected TS-CTB-cre. Comparatively, almost no TS-STB became fusion(+) when co-cultured with *Tg-*infected TS-CTB-cre. Instead, nearly all TS-STB were still exclusively red (fusion(-)) at 48 hours of co-culture (**Figure 1D, S1B**). This result convincingly shows that *Tg-*infected TS-CTB almost completely lose the ability to fuse with TS-STB.

Due to the current unavailability of strongly blue or far-red fluorescing *T*. *gondii* for live imaging experiments, representative images of each condition are shown with IF-staining of *T. gondii* (**Figure S1C**). Live/dead staining (using Trypan blue) of mock-infected and *Tg-*infected TS-CTB-cre indicate that there is no statistically significant difference between the viability of mock-infected cells and *Tg-*infected cells in 3-D culture at 24, 48, or 72 hpi (**Figure S1D**), meaning the robust arrest in fusion is not due to overwhelming levels of cell death of *Tg-*infected TS-CTB-cre.

### Bulk RNA-seq and morphological assessment of *Toxoplasma gondii* infected TS-CTB suggests infection-induced cell fate disruption

Our group recently showed that when TS-CTBs are induced to develop into TS-STB they recapitulate the *T. gondii* resistance phenotype of placental explants and primary trophoblasts^6,8^. In that study, we also generated RNA-seq data from mock-infected and *Tg-*infected TS-CTB that is used in this study for independent analysis of cell differentiation signatures that might be induced by infection. We included transcripts with a log2fold change of >1.5 and p-value of < 0.05 during infection (n = 154 transcripts) in gene ontology analysis. Significantly enriched GO terms during *Tg-*infection include multiple cell differentiation-associated pathways like *Extracellular matrix organization* and *Tissue development.* (**Figure 2A**). Such strong representation of developmental pathways during *T. gondii* infection is unusual and prompted further investigation. These results, particularly in a stemmed cell type like TS-CTB, might indicate differentiation induction during parasite infection. Full output of GO terms also includes pathways that are more typical of infection datasets like *cytokine-mediated signaling pathway* and *cellular response to organic substance* (**Figure S2A**). Gene set enrichment analysis^41^ (GSEA) of the same dataset robustly identified *Epithelial to mesenchymal transition* (EMT) as the most significantly enriched gene set in *Tg-*infected CTB (**Figure 2B**). Transcript increases contributing to this result include multiple matrix metalloproteases (*MMP1, MMP3*), cytoskeletal proteins (*VIM*, *STC1, ACTA2*), collagens (*COL3A1, COL6A3*), and EMT transcription factors (*PRRX1, ZEB1, ZEB2, TWIST1, SNAI2*) (**Figure S2B**). EMT is an essential process underlying CTB differentiation to EVT, suggesting transcriptional changes induced by *Toxoplasma* infection could result in developmental transition towards EVT^42^, thus antagonizing infection-resistant STB formation. We also detected modest but significant changes in known fate markers and transcriptional modulators specifically related to STB and EVT identity including induction of *NRG1, ASCL2,* and *MMP9* along with reduction of *MSX2, ERVW-1, ERVFRD-1,* and *GCM1* (**Figure S2B**). These data suggest that *Tg-*infection polarizes TS-CTB away from a STB-like fate by inducing an EVT-like fate after just 24 hours of infection.

**Figure 2:**
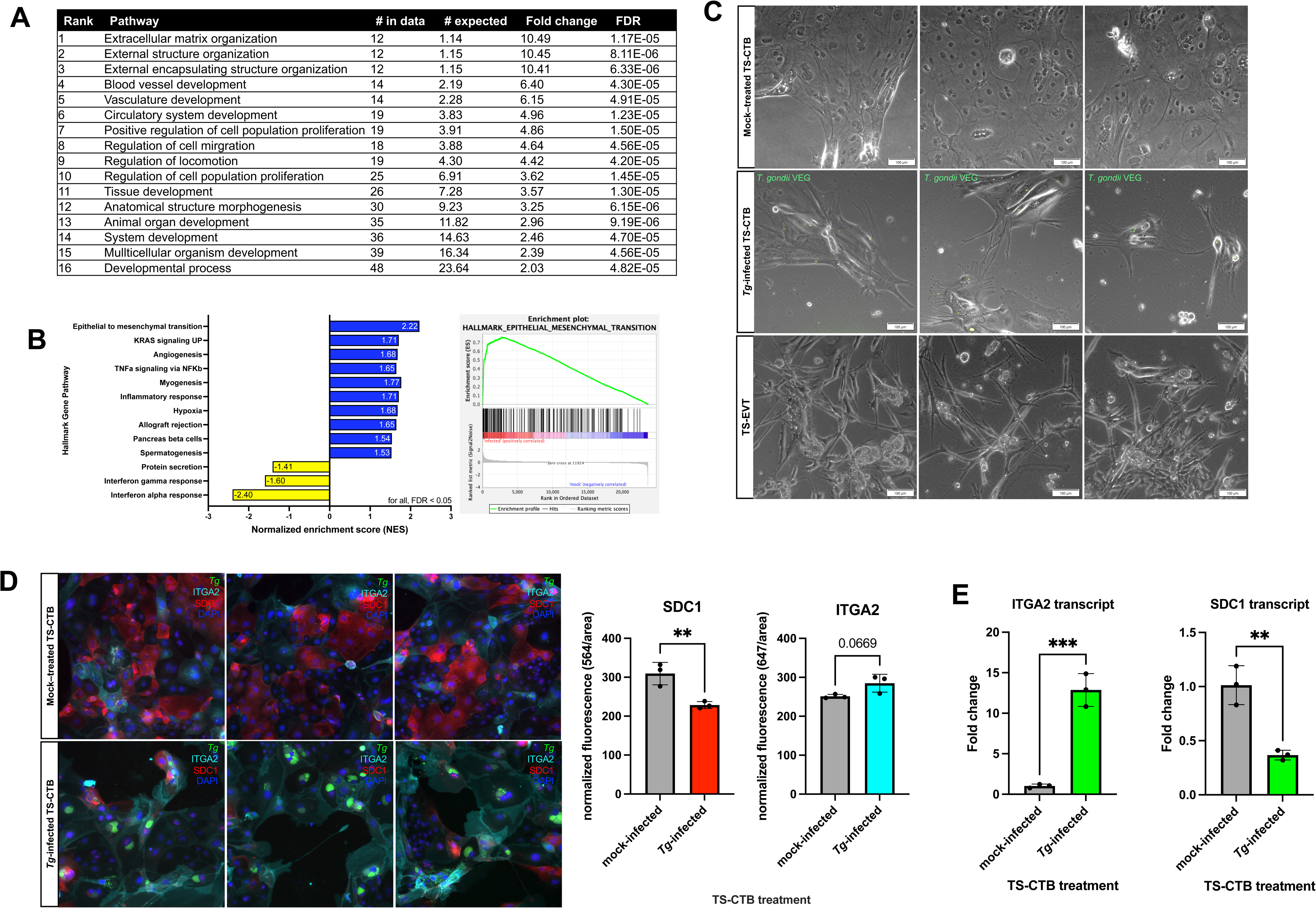
*T. gondii* infection of TS-CTB initiates EVT-like morphology and transcript expression. (A) Pathways from GO term analysis of transcripts with log2fold enrichment > 1.5 and p < 0.05 between *Tg-*infected and mock-infected TS-CTB included in analysis (∼154 gene transcripts). Top 16 pathways ordered by lowest FDR were selected and re-ranked by fold change. (B) Gene set enrichment analysis of all significantly differentially expressed transcripts between mock-infected and *Tg-*infected TS-CTB from bulk RNA-seq of a 24 hour infection. (C) Qualitative cell morphology analysis of mock-infected and *Tg-*infected TS-CTB, and fully differentiated chemically-induced TS-EVT. TS-CTB mock- and *Tg-* infections conducted in TS basal medium + Y27632 for 48 hours. Images taken pre-fixation at 10x magnification. (D) Immunofluorescent assay of 48 hour mock- or *Tg-*infected TS-CTB. *ITGA2* pseudo-colored cyan for contrast. Points represent 2-3 fields of view measured per biological replicate. Fluorescence quantified by integrated density of the given emission channel (564 for SDC1 and 647 for ITGA2) normalized to measured cell area. Significance calculated by one-sample two-tailed unpaired t-test (** p < 0.01). (E) qPCR of *ITGA2* and *SDC1* of 48-hour mock-infected versus *Tg-*infected TS-CTB. GAPDH used as housekeeping gene. Points each summarize three technical replicates of one biological replicate. One-sample two-tailed unpaired t-test performed on linear ΔCT values and results for significance projected onto fold change (** p < 0.01, *** p < 0.001).

Qualitative morphological assessments of *Tg-*infected compared to mock-treated TS-CTB indicate significant morphological changes from typical appearance (**Figure 2C**). Infected cells gain processes and extensions while losing tight cell-cell junctions and other hallmarks of epithelial morphology. The invasive, motile appearance of *Tg-*infected TS-CTB is reminiscent of TS-EVT (**Figure 2C**). Morphology changes appear dependent on the cell density and extracellular space availability as densely plated cells remain morphologically unchanged and do not have altered motility as determined by scratch assay (**Figure S2C**). In combination, the transcriptional and morphological changes observed specifically during infection with *Toxoplasma gondii* suggest a departure from epithelial CTB identity.

To understand if infection-induced morphology changes are a result of cell fate disruption we assessed the levels of *ITGA2* and *SDC1* after infection. *ITGA2* has been consistently identified as a marker of cell column cytotrophoblasts (CCC) that immediately precede differentiation to EVT^17,15^. Oppositely, SDC1 is a marker of STB identity across nearly all levels of maturity^29,15^. The changes in these markers help to identify the directionality of early fate changes in differentiating TS-CTB, with *ITGA2* indicating EVT development and *SDC1* indicating STB development. When we quantified *ITGA2* and *SDC1* in *Tg-*infected versus mock-treated cells, we observed strong enrichment of *ITGA2* and reduction of *SDC1* after 48 hours of *Tg-*infection (**Figure 2E).** Similar results were observed when using a Type II *T. gondii* strain (ME49), though the magnitude of *ITGA2* and *SDC1* changes is lower than for the RH strain used in all other experiments (**Figure S1F**). Quantitative immunofluorescent staining shows a significant reduction of SDC1 and a modest, yet not significant, increase in ITGA2 at the protein level (**Figure 1F**). The simultaneous induction of *ITGA2* and reduction of *SDC1* suggests that *Tg-*infection predisposes TS-CTB to an EVT-like fate in opposition of an STB-like fate.

We also used this transcriptional assay to examine the role of known secreted *T. gondii* proteins in driving this response in TS-CTB. We found that TgMYR1^43,44^, TgWIP^21^ and TgROP47^45^ knockouts all still significantly induced *ITGA2* and reduced *SDC1*, indicating that these parasite genes are not necessary for infection-induced fate disruption in TS-CTB.

### ScRNA-seq reveals coordinated divergence of *Tg-*infected TS-CTB away from STB fate and toward EVT fate

To interrogate infection-mediated changes in cell fate with higher resolution, we sequenced trophoblast stem cells using scRNA-seq. TS-CTB and *Tg-*infected TS-CTB samples were cultured for the duration of infection in basal medium, while TS-EVT and TS-STB were fully differentiated for 8 and 6 days respectively using previously published differentiation culture conditions^29^. Using combinatorial barcoding to uniquely label the cells without the use of microfluidics (that would preclude inclusion of highly fused TS-STB), 10,662 individual cells were sequenced, and from them, six distinct clusters identified (**Figure 3A**). Cells used for sequencing exhibited morphological changes consistent with indicators of the target trophoblast cell type (**Figure S3A**). This design allows us to draw a putative developmental trajectory within these samples (**Figure S3B**) that was confirmed by binning cells based on biological sample origin, marker expression, and pseudotime analysis (**Figure 3B, 3C, S3B**). The developmental trajectory includes three canonical clusters with TS-STB and TS-EVT representing the most terminally differentiated cells and TS-CTB representing bipotent trophoblast stem cells. We identify three clusters of intermediate cell fate which we have named: *proto TS-STB 1*, *proto TS-STB 2*, and *TS-EVT predisposed* (**Figure 3A**). Using *CGA, EGFR,* and *TEAD1*, which are each highly expressed STB, CTB, and EVT markers, respectively, we can see the progression of differentiation across clusters (**Figure 3C)**. *CGA* in particular is a robust indicator of trophoblast identity progression as its expression goes from minimal in EVT to moderate in CTB to very high in STB. TS-STB are defined by high levels of *CGA, PAPPA, CGB1/3/5/7, KISS1, SDC1,* and *CYP19A1. Proto TS-STB 1* and *2* intermediately express these same markers and highly express pre-STB fusion markers like *ERVW-1* and *ERVFRD-1.* TS-EVT are defined by high levels of *TEAD1, TCF7L2, GCM1, FN1, MMP2, HLA-G, HTRA4, DIO2, ITGA5,* and *NOTUM* (**Figure 3B**). The TS-CTB cluster contains cells uniquely expressing *TFAP2C, SP6,* and *LYN*, although this cluster shares many of its markers with the *TS-EVT predisposed* cluster. *TS-EVT predisposed* cells express typical stemness markers *TEAD4, YAP1, LRP2,* and *TP63* that are shared with the TS-CTB cluster. However, cells in the *TS-EVT predisposed* cluster simultaneously strongly express transcripts that indicate they may be predisposed against an STB fate and towards an EVT fate, including *CDH1, NOTCH1, ITGB4, BCAT1, MKI67, TEAD1,* and *TCF7L2*. These cells also express comparatively less *CGA* than cells of the canonical *TS-CTB* cluster, suggesting they are further from an STB identity (**Figure 3B, 3C)**. We believe that the *proto TS-STB 2* fate represents the underlying predisposition of TS-CTB to fuse to become TS-STB-like in basal + Y27632 media. Cells in this cluster come from all biological samples (**Figure S3B**) and may broadly capture cells unable to respond to chemical induction.

**Figure 3:**
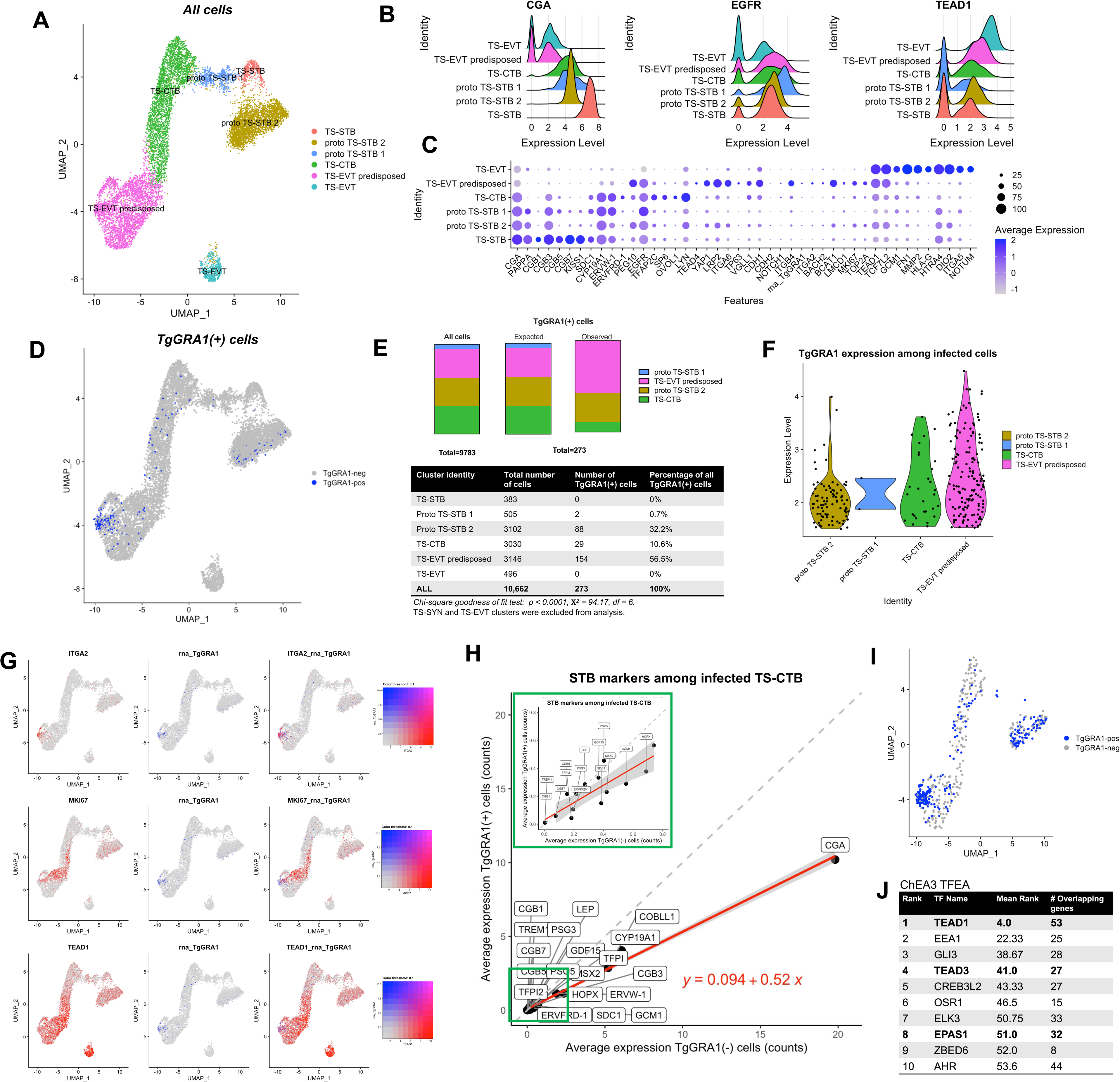
scRNA-seq reveals coordinated development of *Tg-*infected TS-CTB towards an EVT-predisposed fate. (A) UMAP of all sequenced cells with cluster names determined by compiling multiple analyses to determine cell types using marker expression, biological sample analysis, and pseudotime (filtering cutoffs used: RNA features < 3500, RNA counts < 15000, % mitochondrial reads < 25). (B) Ridge plots of STB (CGA), CTB (EGFR), and EVT (TEAD1) markers across clusters (C) More marker transcript expression across clusters. Transcripts were selected from datasets analyzing both *in vitro* and *ex vivo* primary placental trophoblasts. (D) TgGRA1(+) cells among all cells. Cutoff for TgGRA1 expression > 1.5. (E) Proportional representation of TgGRA1(+) cells per cluster. Chi-square goodness of fit test to determine significance of difference between observed and expected distribution of TgGRA1(+) cells across clusters. TS-STB and TS-EVT clusters excluded from this analysis as these mature cells should only be generated through chemical induction in samples that will not experience infection. p < 0.0001, X^2^ = 94.17 on df = 6. (F) TgGRA1 transcript expression among TgGRA1(+) cells across clusters (G) Co-visualization of proto-EVT markers with TgGRA1(+) cells. Color according to scale indicates simultaneous expression level of each transcript. (H) Average counts for a given STB marker transcripts in TgGRA1(-) cells versus TgGRA1(+) cells. Slope of the linear regression of these points is a proxy for level of global reduction or induction of STB transcripts between cell populations with slope > 1 indicating induction of STB transcripts in TgGRA1(+) cells and slope < 1 indicating reduction of STB transcripts in TgGRA1(+) cells. Slope = 0.52 with f-statistic 2259 on 1 and 18 DF, p < 2.2e-16). (I) Down-sampled dataset with equal numbers of TgGRA1(+) and TgGRA1(-) cells. (J) Results of ChEA3 transcription factor enrichment analysis tool of 114 significantly increased transcripts between TgGRA1(+) and TgGRA1(-) cells. TFs are ordered according to their mean rank across all the compiled datasets in ChEA3 tool. Overlapping genes indicates the number of genes in the submitted list that are part of the total list of interactors for a given TF.

Due to *Toxoplasma* being an intracellular eukaryote, scRNA-seq of parasite-infected cells offers the unique opportunity to capture both host and pathogen transcripts simultaneously. Using a transcript that is exclusively and highly expressed by *T. gondii* called TgGRA1, we identified cells with an internal parasite from the infected TS-CTB biological sample (**Figure 3D**). Discrimination using *TgGRA1* allows us to distinguish between directly infected and bystander cells in the infected TS-CTB biological sample (**Figure S3C**), while the use of only a single *Toxoplasma* gene ensures that the clustering of trophoblast cell types is not altered. After excluding TgGRA1(+) cells from biological samples that were not infected (putative doublets with mock-treated TS-CTB, TS-STB, and TS-EVT), we were left with 273 *TgGRA1*(+) cells (**Figure 3D**). We posited that if there were no impact of infection on cell identity, that *TgGRA1*(+) cells would be distributed equally across the four intermediate fate cell type clusters. Instead, we found that the majority (56.5%, 154/273) of *TgGRA1*(+) cells localized to the putative TS-EVT predisposed cluster, indicating coordinated cell identity departure among infected TS-CTB (**Figure 3E**). The aggregation of TgGRA1(+) cells in the *TS-EVT predisposed cluster* is significant as determined by chi-square test (p-value < 0.0001) (**Figure 3E**). The top and bottom 10 most differentially expressed transcripts of TgGRA1(+) cells compared to all TgGRA1(-) cells include transcripts related to both early EVT and STB identity respectively (**Figure S3D**).

*TgGRA1* transcript can be used as a proxy for the number of parasites inside a given cell, with higher levels of *TgGRA1* correlating with robustly replicating parasites inside full parasitophorous vacuoles. Notably, the mean value of *TgGRA1* transcript is highest among TgGRA1(+) cells in the TS-EVT predisposed cluster (**Figure 3F**). This suggests that, compared to TgGRA1(+) cells in other clusters, there is a larger number of parasites per cell in infected TS-EVT predisposed cells compared to infected cells in other clusters. Therefore, the observed cell fate changes are happening most often in response to a robust and progressive *Toxoplasma* infection.

To delineate early TS-EVT progression, we have highlighted *ITGA2, MKI67,* and *TEAD1* to represent progressive stages of EVT precursor identity. *In vivo,* these markers correlate to the progression of proliferative *ITGA2*(+) cell column cytotrophoblasts (CCCs) to differentiated EVT that will invade the maternal decidua^15,17^. These transcripts are co-visualized with *TgGRA1* to indicate the relative position of *TgGRA1*(+) cells at the beginning of this trajectory (**Figure 3G**). This selective localization of TgGRA1(+) cells suggests that *Tg-*infected TS-CTB transcriptionally resemble early proliferative EVT precursors while being highly distinct from TS-derived STB.

To assess global transcriptomic changes related to TS-STB fate, we plotted STB marker expression between downsampled TgGRA1(+) cells versus TgGRA1(-) cells within the infected TS-CTB sample (**Figure 3I)**. We then performed linear regression among these points and calculated a slope of 0.52 (F-statistic 2259 on 1 and 18 DF, p-value < 2.2e-16), suggesting a global reduction of the STB developmental program during *Toxoplasma* infection (**Figure 3H**).

To identify transcription factors that could be implicated in *Tg*-induced developmental changes we conducted transcription factor enrichment analysis (TFEA) with ChEA3^46^. Among the 114 transcripts significantly (p < 0.05) increased between TgGRA1(+) and TgGRA1(-) cells TFEA identified TEAD1 (#1), TEAD3 (#4), and EPAS1 (#8) in the top 10 rank-ordered TFs (**Figure 3J**). Importantly, each of these TFs has been implicated as a major regulator of EVT developmental transition^47^, further supporting a fate change in infected TS-CTB that is antagonist to STB formation.

Notably, with this sequencing approach we also capture the transcriptional profiles of *Toxoplasma* parasites. The transcript differences driving the two resolved parasite clusters are modest with only 12 significant differentially expressed transcripts between clusters (p < 0.05) (**Figure S3E-F**). The lack of significant differences is in step with the idea that *Toxoplasma* largely does not change its transcriptome according to the cell type it infects.

With this approach, we observed a universal reduction of STB markers and induction of EVT markers in TgGRA1(+) versus TgGRA1(-) cells. TgGRA1(+) cells cluster directionally with TS-EVT predisposed cells, indicating that *T. gondii* infection drives transcriptional polarization towards EVT lineage in stem-state trophoblasts.

### Multiomic snRNA-seq and snATAC-seq reveal shared chromatin accessibility profiles between *Tg-*infected TS-CTB and early TS-EVT

Epigenetic regulation has been increasingly implicated as a principal driver of developmental programming in diverse cell types, including placental trophoblasts. To investigate the epigenetic landscape of *Toxoplasma*-infected TS-CTB compared to their differentiated counterparts, we sequenced 43,822 single nuclei with paired snRNA-seq and snATAC-seq. *Tg-* infected TS-CTB, mock-infected TS-CTB, day 4-differentiated TS-EVT (D4 TS-EVT), and day 8-differentiated TS-EVT (D8 TS-EVT) samples were sequenced. Though we initially intended to estimate the population of TgGRA1(+) cells by marker expression informed by our previous single-cell dataset, we were surprised to detect non-random expression of *TgGRA1* in the infected TS-CTB sample. This putative *TgGRA1* expression is almost directly in-line with the estimation of TgGRA1(+) cells that we would have made according to marker expression cutoffs. If parasite nuclei were NOT host-nuclei associated, we would also expect that they would both fail quality control filtering cutoffs for human cells and/or that they would localize randomly throughout the UMAP, which they do not (**Figure 4C**). There is also not enough sequence homology between *TgGRA1* and human transcripts to support it being mistakenly identified as a different non-randomly expressed human transcript. These results all point to the association between host cell nucleus and TgGRA1 transcripts being maintained through the single-nucleus microfluidic prep. In an excess of caution, the majority of our bioinformatic analyses have been performed on and visualized according to biological sample-level or cluster-level identities (**Figures 4A, 4B, 4D 4F-H**) which are unaffected by TgGRA1(+/-) status.

**Figure 4:**
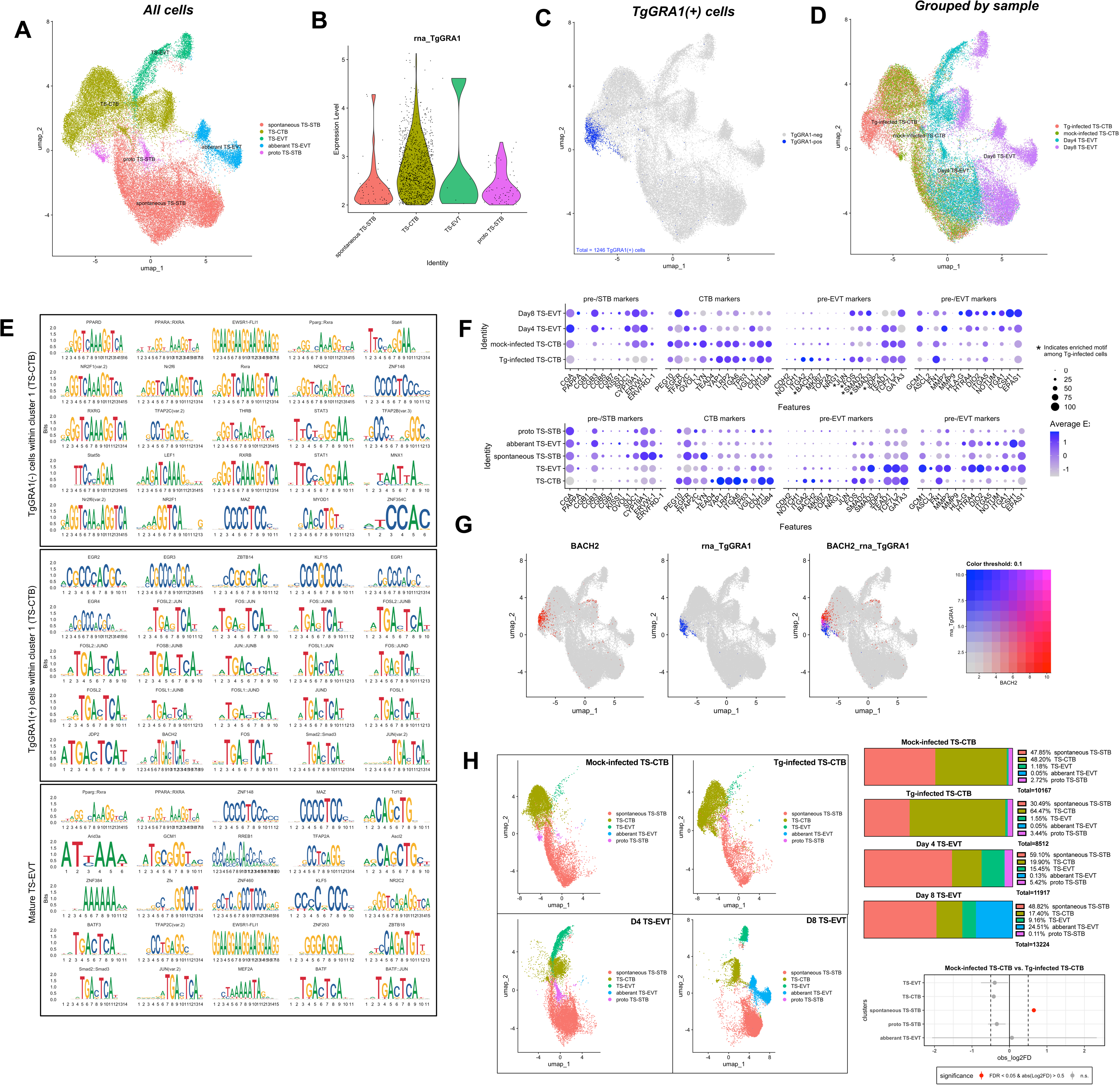
snRNA-seq and snATAC-seq reveal shared epigenetic signatures between Tg-infected TS-CTB and TS-EVT. (A) UMAP of all sequenced cells with cluster names determined by compiling multiple analyses to determine cell types using marker expression, biological sample analysis, and pseudotime (B) TgGRA1 expression among intermediate fate clusters. (C) TgGRA1(+) cells distributed on UMAP. (D) UMAP coded by biological sample origin. (E) Motif enrichment analysis among differentially expressed chromatin accessibility peaks between mature TS-EVT, TgGRA1(+) cells within TS-CTB cluster, and TgGRA1(-) cells within TS-CTB cluster. Top 25 differentially motifs among differentially accessible peaks for each sample (F) Marker transcript expression among biological samples. *indicates enriched motif among *Tg-*infected cells. (G) BACH2 transcript expression co-visualized with TgGRA1 expression. (H) Distribution of cells from each biological sample origin into cluster identities. Permutation test to determine if the distribution of cells across clusters between mock- and *Tg-*infected TS-CTB biological origin sample is significantly different, n = 10,000 permutations.

Firstly, the snRNA-seq data replicates many of our original findings from the prior scRNA-seq dataset (**Figure 3**). Repeated findings include the dense aggregation of 1246 TgGRA1(+) cells at one particular region of the TS-CTB cluster, again indicating coordinated cell fate between *Toxoplasma*-infected cells (**Figure 4C**). We also again observe the highest level of TgGRA1 transcript in the TS-CTB cluster. (**Figure 4B**). We find similar markers expressed by Day 4- and Day 8-differentiated TS-EVT as those expressed by previous TS-EVT, with greater nuance introduced by temporally separated samples (**Figure 4F**). But most importantly, we continue to observe spontaneous TS-STB formation through all biological samples. These spontaneous TS-STB populate the largest cell cluster and result in variable STB marker expression throughout most biological samples (**Figure 4A, 4D, 4F**). We continue to take advantage of this predisposed fate as a means to provide the counterbalance STB developmental identity that would normally be precluded by use of sn-sequencing.

Using snATAC-seq data, motif enrichment analysis reveals vastly different TF binding sequences enriched in differentially accessible regions in *Tg-*infected compared to mock-infected TS-CTB (**Figure 4E**). Binding motifs enriched among differentially accessible peaks in mock-infected TS-CTB include those recognized by PPARA/D/G complexes with and without RXRA/G. We also see enrichment of motifs recognized by canonical CTB stemness regulators LEF1 (TCF4) and TFAP2C/B. Several STATs are also represented including STAT1/3/5B. Among *Tg-* infected TS-CTB, we see an entirely different set of enriched binding motifs. These include recognition sites of known *Toxoplasma* induced TFs including EGR1/2/3/4. We also see extensive representation of a TGA_TCA motif that can be used by various homo- and heterodimeric combinations of FOS/L1/L2 and JUN/B/D (**Figure 4E**). The same motif is recognized by BACH2 and SMAD2:SMAD3, both of which regulate cell development in trophoblastic and non-trophoblastic contexts^48,49^. BACH2 is also enriched in transcript expression among *Tg-*infected TS-CTB (**Figure 4F, 4G, S4A**). Finally, in TS-EVT, we found representation of canonical EVT-regulating TFs including GCM1 and ASCL2^50,28^. We also detected enrichment of the same TGA_TCA motif (for SMAD2:SMAD3, JUN, BATF and BATF:JUN) in these TS-EVT, suggesting TF recognition of this sequence could underlie both the transcriptional drivers of infection-induced TS-CTB development and canonical EVT development.

Among motifs represented within top differentially accessible peaks, the transcription factor BACH2 represents a particularly interesting candidate as both its transcript and chromatin accessibility are enriched in *Tg-*infected TS-CTB (**Figure 4G, S4A)**. Described previously for its role in T-cell differentiation, BACH2 expression has also recently been found to be uniquely enriched in cell column cytotrophoblasts (CCC), or, EVT precursors^15^. We see higher accessibility of the BACH2 promoter in addition to several exons and intergenic regions in *Tg-*infected compared to mock-infected TS-CTB (**Figure S4A**). There are 9 linked peaks across the genomic locus, indicating that chromatin accessibility increases across the entire region in a coordinated fashion during *Toxoplasma* infection (**Figure S4A**). Motif footprint plots indicate both the rate of Tn5 insertion immediately upstream and downstream of a given TF binding site motif compared to at the motif itself. Here, we see increased accessibility immediately upstream and downstream of BACH2 motifs in TgGRA1(+) TS-CTB compared to TgGRA1(-) TS-CTB (**Figure S4B**). We also see high occupancy of the motif by its client TF (or other TFs capable of recognizing the same sequence), indicated by the precipitous drop in Tn5 insertions at the motif sequence (**Figure S4B**). Together, these data indicate several shared epigenetic characteristics between *Tg-* infected TS-CTB and TS-EVT, supporting the hypothesis that *Toxoplasma* infection induces cell fate changes that propagate EVT development to antagonize STB development.

Finally, based on synthesis of many datasets of TS cells, we generally expect ∼50% (or more) of cells in any given condition to transcriptionally resemble spontaneously fused TS-STB-like cells. This is observed in the mock-infected TS-CTB, Day 4 TS-EVT, and Day 8 TS-EVT samples with 48.20%, 59.10%, and 48.82% respectively clustering as spontaneous TS-STB (**Figure 4H**). In contrast, in the *Tg-*infected biological sample we see a statistically significant depletion of spontaneously formed TS-STB with only 30.49% of cells mapping to this cluster (FDR < 0.05, and |log2FC > 0.5|, permutation test; n=10,000, method as in ^51^) (**Figure 4H**). Instead, the TS-CTB cluster gains representation during *Toxoplasma* infection, growing from only 48.20% in the mock-infected condition to 64.47% in the *Tg-*infected condition. This redistribution of cell types between mock- and *Tg-*infected TS-CTB continues to emphasize how parasite infection specifically antagonizes spontaneous TS-STB formation.

### *Toxoplasma*-induced developmental changes are detectable in *ex vivo-*infected chorionic villi

Given the clear similarities in the response of TS-CTB to *T. gondii*, we wanted to determine if similar transcriptional changes would occur in placental villous explants. To date, achieving robust *ex vivo* infection of trophoblasts in late-gestation chorionic villi (CV) with *Toxoplasma* has been challenging. Previous experiments with late-gestation villi suffer from low parasite invasion rate as the STB layer is strongly resistant to *Tg-*infection^6,7,8^, a phenotype that likely intensifies with gestational age. To increase parasite infection in the CTB layer, we first performed serial blunt transverse slicing of the villi. We then infected sliced villi with 10 million parasites for 48 hours (**Figure 5A**) and processed them for RNA-seq.

**Figure 5:**
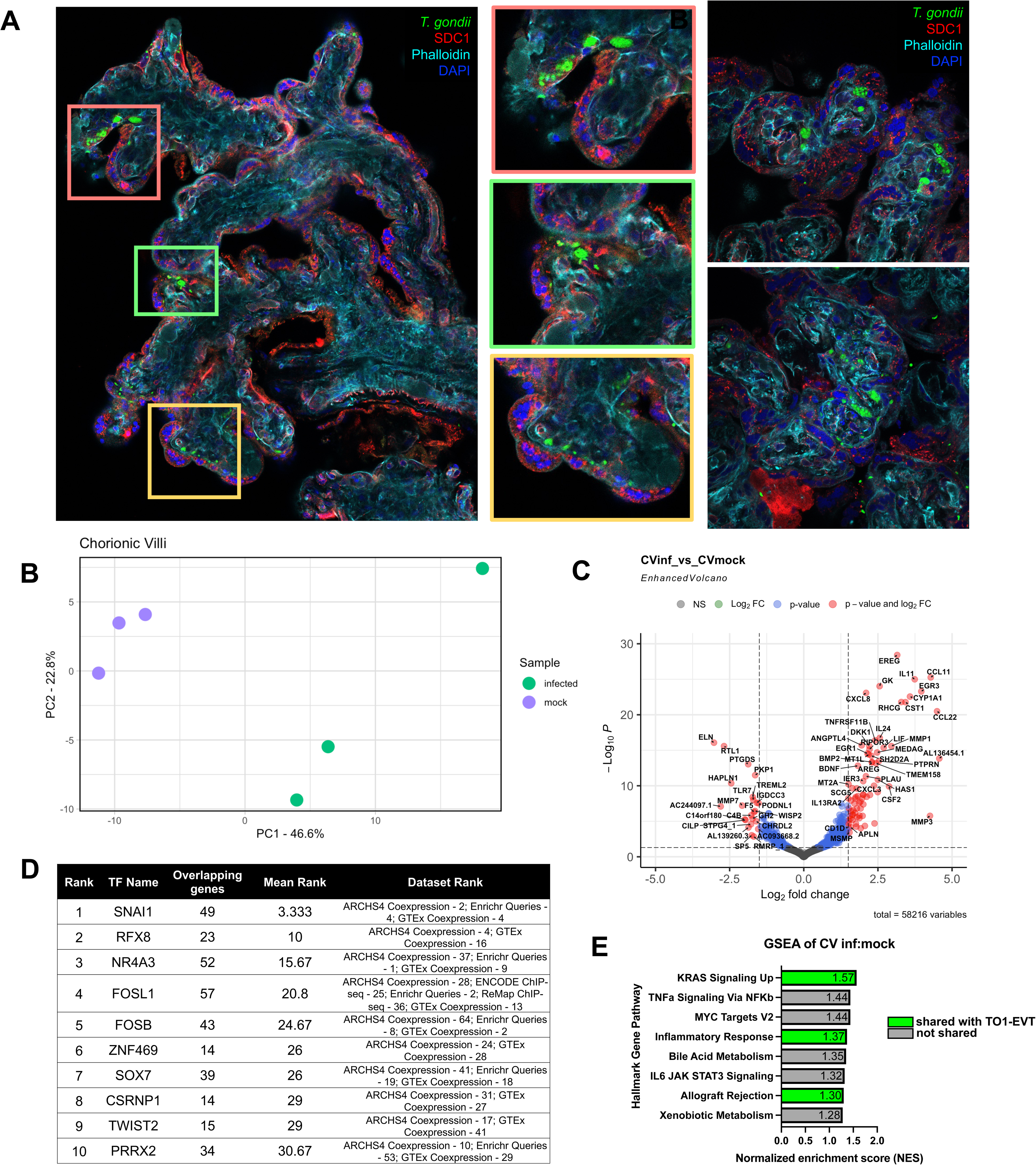
Developmental disruption is detectable in *Tg*-infected chorionic villi. (A) IF image of dissected, sliced *Tg-*infected chorionic villus. Center panels represent regions of high infection. Right panels show 60X zoom of intracellular replicating *T. gondii*. (B) PCA of mock versus infected chorionic villi. (C) Differential transcript expression between mock- and *Tg-*infected chorionic villi. (D) ChEA3 TFEA of differentially enriched transcripts in *Tg-*infected chorionic villi (E) GSEA output of *Tg-*infected versus mock-infected chorionic villi, all hallmark pathways included with p < 0.05.

Bulk sequencing of *Tg-*infected chorionic villi indicates enrichment of many of the same *Tg-*induced transcripts observed in other models. *Tg-*infected and mock-infected samples separate appropriately in PCA (**Figure 5B**). *Tg-*enriched genes conserved between models include *MMP1/3*, *EGR1/3*, and *DKK1* (**Figure 5C**). Important *Tg-*enriched genes from previous work with primary trophoblasts were also detected in *Tg*-infected CV including *CCL22, IL11, CCL11,* and *IL24*^,52^. Several of these transcripts have been absent from analyses of other models in this study, which may be due to the natural immunological variation between models and the heterogeneous cell types that they consist of (TS cells, TOs, and CV). This analysis also detects *Tg-*enriched transcripts that have been directly implicated in both human and mouse placental differentiation like *BMP2*^53,54^. Additionally, we detect strong induction (L2FC = 2.88, rank in dataset ordered by L2FC = 23) of *CSF2*, a uNK cell-derived cytokine that has recently been shown to potently promote EVT formation in trophoblast organoids^16^.

From this same dataset, TFEA strongly implicates SNAI1 as a driver of the transcripts differentially expressed in *Tg-*infected chorionic villi (**Figure 5D**). Notably, SNAI1 is a canonical driver of EMT and has also been described as dictating EVT transition^47^. Though the *Hallmark EMT* pathway is not significantly enriched in *Tg-*infected villi by GSEA (**Figure S5A**), we note three shared enriched hallmark pathways between infected villi and EVT-differentiated trophoblast organiods (**Figure 5E**, **Figure 6**): *KRAS Signaling Up*, *Inflammatory Response*, and *Allograft Rejection* (**Figure 5E**). Despite that the structure of chorionic villi makes it difficult to achieve robust parasite infection *ex vivo*, we see important retained transcriptional signatures of *Toxoplasma* infection that suggest developmental disruption is conserved at the tissue level.

**Figure 6:**
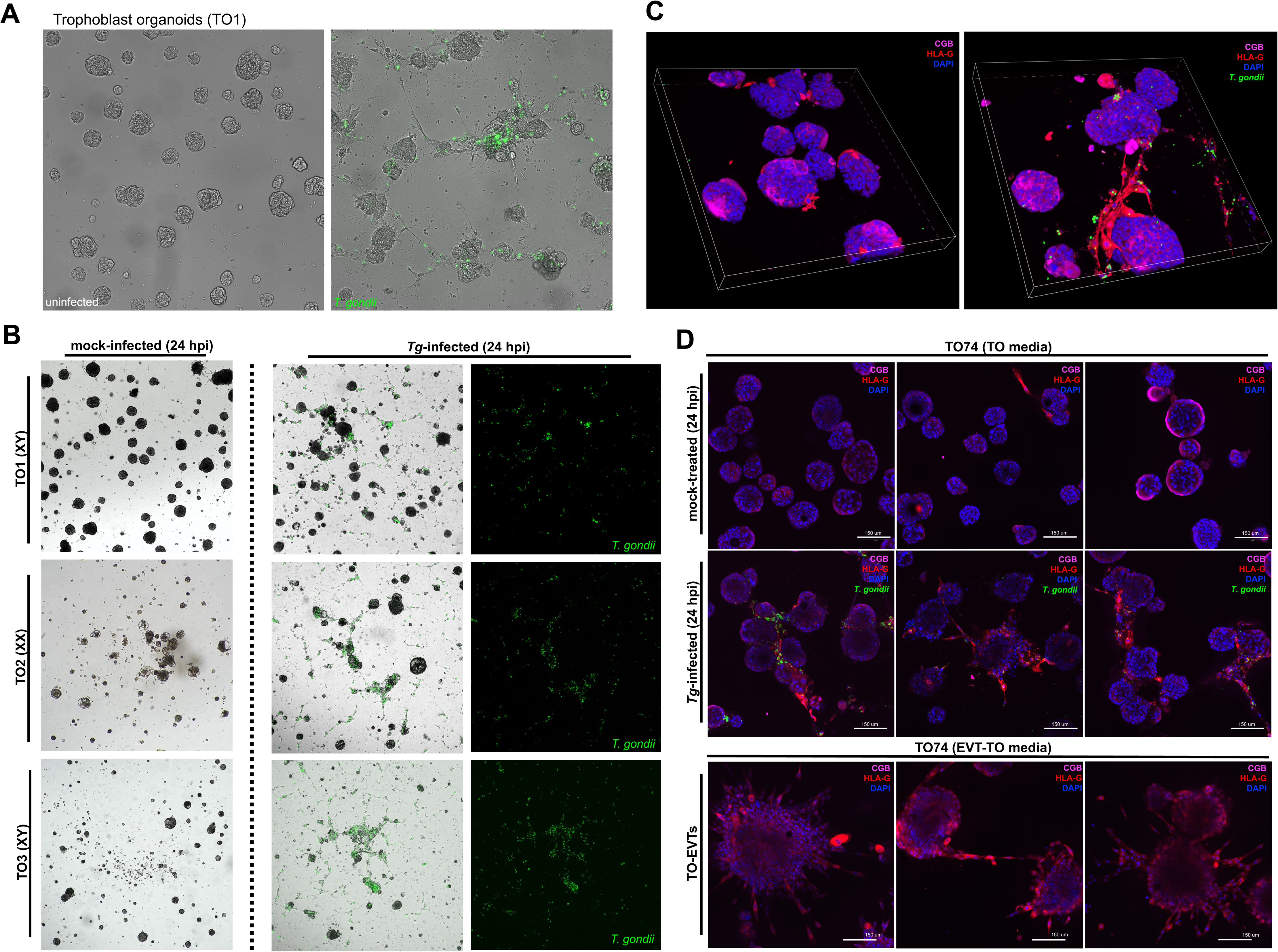
Trophoblast organoids develop HLA-G(+) EVT upon infection with *T. gondii*. (A) Representative images of unfixed uninfected or *Tg-*infected third trimester-derived trophoblast organoids (TOs) 24 hours post infection. Organoids cultured and infected on Matrigel layers in 8-well chamber slides. (B) Representative images of paired unfixed mock- and *Tg-*infected TO1, TO2, and TO3. Green-fluorescent *T. gondii* shown for *Tg-*infected TOs (C) and (D) 3D and 2D confocal immunofluorescent imaging of mock-infected and *Tg-*infected TO1 showing expansion of HLA-G positive cell population in *Tg-*infected TOs compared to mock-infected TOs and TO1-EVTs.

### Trophoblast organoids develop HLA-G positive EVT during *Toxoplasma gondii* infection

Growing evidence suggests that signaling cues between placental cell populations is essential to the deployment of developmental processes^14,16,55,56^. The trophoblast organoids used in this study (originally derived in ^57^) allow all three trophoblast types (CTB, EVT, and STB) to be cultured simultaneously and therefore may recover inter-cell type signaling cues. TOs are propagated and cultured in Matrigel domes and grow both CTB and STB in an “inside-out” conformation, where mononuclear CK19+/SDC-CTB are on the outside of a fused CK19+/SDC1+ STB core (**Figure S6A**). Minimal spontaneous EVT formation occurs in normal TO culture, rather EVT induction requires temporal supplementation and removal of exogenous NRG1 in the culture media over the course of several weeks^58,59,60^. We infected TOs with *T. gondii* to understand how infection-induced trophoblast development might manifest in a multicellular organoid model.

Strikingly, just 24 hours of *T. gondii* infection in TOs was sufficient to induce HLA-G(+) cellular outgrowths that morphologically resemble chemically induced TO1-EVTs (**Figure 6A-D**). HLA-G(+) cells resulting from *Tg-*infection were observed across three genetically distinct TO lines derived from the placentas of both male (XY) and female (XX) fetuses (**Figure 6B**). We observed the strongest outgrowth phenotype when TOs were cultured and infected on Matrigel layers in 8-well chamber slides. *Tg-*infected TOs that were infected and maintained in suspension culture did not produce cellular outgrowths but did qualitatively increase HLA-G staining (**Figure S6B**). Similarly to infected TS-CTB, *Tg-*infected TOs exhibited increased *ITGA2* expression and decreased *SDC1* expression compared to mock-infected TOs (**Figure S6C**). We qualitatively observed *T. gondii* primarily in HLA-G(+) regions and suspect that direct infection of the outer layer CTB progenitors leads to the outgrowth of these elongated cells. In z-stack images, infection-induced elongated processes are found on multiple planes of the Matrigel layer, potentially indicating local invasive capacity rather than directional growth on either the liquid or matrix side (**Figure 6C-D**).

In effort to identify transcriptional drivers of HLA-G(+) cell formation, we conducted bulk RNA-seq of TO1-EVT, mock-& *Tg-*infected TO1/TO2 (**Figure S6D**). Significantly enriched transcripts in *Tg-*infected TOs include many of the same transcripts observed in our initial bulk TS-CTB dataset (**Figure 7C, S6E**) including *EGR1/2/3/4, VIM, GREM1, THY1, LUM, STC1,* and *DCN* (TO1 in **Figure 7C**, TO2 in **Figure S6E**). As we originally observed in TS-CTB, a consistent set of well-expressed extracellular matrix remodeling genes are induced by infection including *MMP1, MMP3, TNC, LUM, COL6A3, COL6A2,* and *COL1A2* (TOs in **Figure S6E, 7C,** TS-CTB reproduced in **S6E**). The similarity between genes induced in *Tg-*infected TOs compared to TS-CTB suggests conservation in the transcriptional programs being impacted by *Toxoplasma*. Surprisingly, we do not observe many transcripts with coordinated enrichment/diminishment patterns between chemically induced TO1-EVT and *Tg-*infected TO1. TO1-EVT do, however, highly express transcripts related to EVT differentiation including Type I interferon-stimulated genes^28^, *MMP2, LAMA4,* and *DIO2* (**Figure 7C**). These results summarize the transcriptional complexity of EVT differentiation in chemically induced TO1-EVT, of which *Tg-*infected TO1 only modestly and partially recapitulate at the resolution of bulk RNA-sequencing.

**Figure 7:**
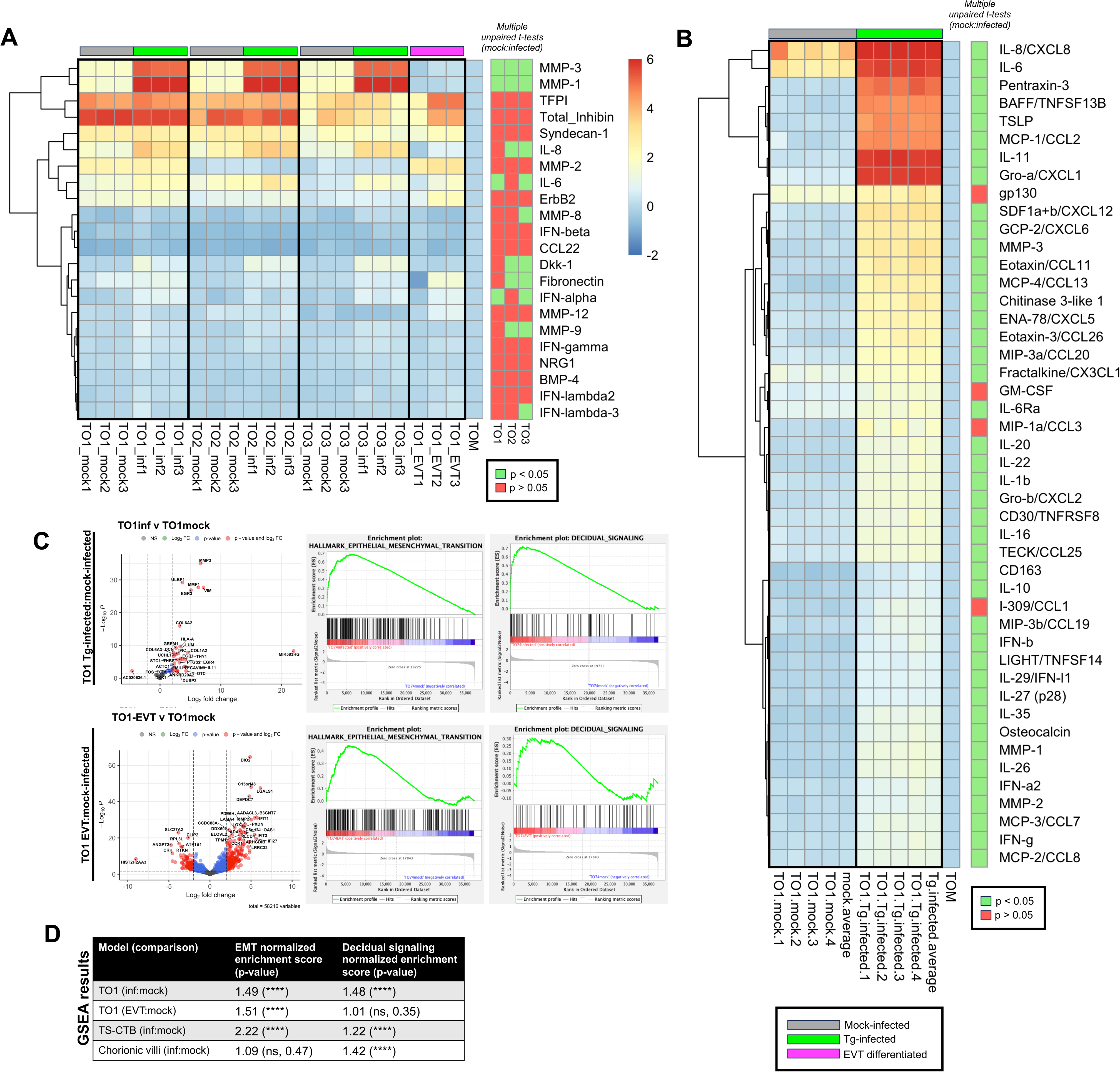
Epithelial to mesenchymal transition and decidual signaling cues induced by *T. gondii* compound to elicit EVT formation across models. (A) Heatmap of 22 analytes from custom Luminex analysis of mock-infected and *Tg-*infected TO1, TO2, and TO3; and TO1-EVT. Data represent log2fold change from TOM (unconditioned media). Multiple unpaired t-tests used to determine statistical significance of the difference between mock-infected and *Tg-*infected samples of each TO line. (B) Same as (A) but data from 46 analytes from mock-infected and *Tg-*infected TO1. 72 total analytes were tested but only those that had a ratio of > 1.5 or < 0.5 between mock- and *Tg-*infected conditions were included in the main figure. All analytes shown in **Figure S8B**. (C) Volcano plots of differentially expressed transcripts from bulk RNA-seq of mock-versus *Tg-*infected TO1, and TO1-EVT versus mock-infected TO1. Corresponding GSEA plots from Hallmark EMT and custom Decidual signaling datasets included. (D) Normalized enrichment scores from GSEA of indicated dataset. P-value significance indicated in paretheses. **** p < 0.0001

GSEA supports the enrichment of epithelial to mesenchymal transition (EMT) in both *Tg-* infected TO1 and TO1-EVT compared to mock-infected TO1 with normalized enrichment scores of 1.49 and 1.51, respectively (p-value < 0.0001) (**Figure 7C-D**). This data is strongly reminiscent of the EMT enrichment we first observed in *Tg-*infected TS-CTB (**Figure 1C, S6E, 7D**), suggesting EMT dictates both canonical and *Tg-*induced EVT lineage specification across models. *Tg-* infected TO1 and TO1-EVT also share other significantly enriched hallmark pathways compared to mock-infected TO1 including *Angiogenesis* and *KRAS Signaling Up* (p-value < 0.05) (**Figure S7A**). Interestingly, *Tg-*infected TO1 also show significant enrichment for *Hypoxia* and *Wnt Beta Catenin Signaling*, pathways that have each been implicated in EVT formation^61,62^. Though these pathways are not enriched during chemical TO1-EVT formation, we find it significant that *Toxoplasma* infection induces developmentally relevant gene expression signatures in progenitor cells. Although infected TOs do not completely recapitulate the transcriptional profile of chemically induced TO1-EVTs, the rapid onset of EVT-related changes in infected TOs supports that *T. gondii* potently activates essential elements of the early EVT developmental program in complete opposition of the STB developmental program

Moving forward, we were keen to understand why infection-induced cell differentiation was more dramatic in trophoblast organoids compared to trophoblast stem cells or chorionic villi. Beyond the ability to recapitulate inter-cell type signaling cues, we hypothesized that a broader repertoire of expressed cytokines could contribute to greater EVT outgrowth. We conducted multi-analyte Luminex analysis of supernatants from mock-infected and *Tg-*infected TO1, TO2, and TO3 (**Figure 7A, 7B**). From this analysis, we detected significant and robust increases in multiple proinflammatory cytokines during *T. gondii* infection including IL-6, IL-8, IL-11, and CCL2 (**Figure 7B)**. This repertoire of cytokines stimulated by *T. gondii* infection of organoids is quite like the repertoire that is expressed by decidual cells. *In vivo,* decidual and placental signaling work in tandem to control the invasion of trophoblasts during placentation^14,16,63,64,65^. Informed by the secretome of decidual organoids^57^, we analyzed the top 15 differentially secreted proteins by uninfected DOs compared to TOs via Luminex. We found that 9 of the 15 canonical DO secreted factors were significantly increased in *Tg-*infected TOs compared to mock-infected TOs (**Figure S7B**). We found no secretion or no significant difference of 5/15 factors and only 1/15 being statistically significantly decreased by *T. gondii* infection **(Figure S7B**).

In exploring the representation of decidua-associated signaling factors across differential expression analyses of our placental datasets, we also found that many are strongly induced by *Toxoplasma* infection at the transcript level (**Figure S7C**). To further quantify the contribution of decidua-associated signaling, we assembled a custom gene set of decidual signaling factors (**Table S8A,** n = 62 transcripts) from both literature and recently published uninfected decidual organoid secretome data^57^. With this gene set, we conducted GSEA across bulk RNA-seq datasets of our *Tg-*infected TS-CTB, TO1, and chorionic villi versus their respective mock-infected counterparts (**Figure S7C**). We find that the decidual signaling gene set is universally and significantly enriched among *Tg-*infected datasets (NES of TS-CTB: 1.22, TO1: 1.48, Chorionic villi: 1.42, all p < 0.0001) (**Figure 7B**). This contrasts with the non-significant enrichment of decidual signaling in the TO1-EVT versus mock-infected (NES: 1.01, p-value = 0.35). The lack of significant decidual signaling gene set enrichment in TO1-EVTs is expected as our data suggests that production of increased decidual signaling cues is a non-trophoblastic process that accelerates EMT-induced EVT outgrowth (**Figure 7C, D**) and, in this case, is specifically induced by *T. gondii* infection. Collectively, *Toxoplasma* infection activates EMT which is then compounded by secreted pro-inflammatory cytokines to entirely misdirect trophoblast lineage specification.

## DISCUSSION

During congenital infection, transmission of the parasite through the placenta exposes trophoblast stem cells to massive biological changes imparted on host cells by *Toxoplasma*. As the placenta must complete the entire process of organogenesis while also functionally supporting the fetus, it is particularly vulnerable to developmental dysregulation. We are aware of only a handful of recent studies that address any element of the connection between trophoblast development and congenital pathogen infection, with these examples being restricted almost entirely to viral systems^11,35,34,66,67^.

In this study, we thoroughly characterize how *Toxoplasma* infection antagonizes STB formation by causing arrest of hormone secretion and cell fusion. Instead, infection induces a unique intermediate and polarized developmental fate of trophoblast stem cells that mechanistically antagonizes the STB developmental program. Our study demonstrates distinct morphological and transcriptional changes that redirect trophoblast stem cell fate away from the parasite-restrictive STB lineage and towards the parasite-permissive EVT lineage. *Tg-*infected TS-CTB resemble a specific niche of EVT precursors that are then functionally prevented from completing differentiation checkpoints, like fusion, to form STB. This is similar to recent publications that show SARS-CoV-2 and Zika viruses cause STB fusion arrest in trophoblast organoids and trophoblast stem cells, although our study improves on these by using a functional fusion assay^34,35^. We observe signatures of these phenotypes across placental models, including trophoblast organoids, trophoblast stem cells, and chorionic villi. We go on to validate *Toxoplasma* infection-induced differentiation in a self-contained multi-cell type organoid model where we observe rapid outgrowth of HLA-G(+), locally invasive EVT. Finally, we implicate epithelial to mesenchymal transition and phenocopied decidua-associated cytokines in supporting infection-induced cell differentiation towards EVT at the cost of STB.

While the direct relevance of developmental dysregulation to placental health and congenital transmission is unknown, it might disproportionately skew trophoblast populations to weaken the innate barrier function of the STB layer. Thorough histopathological data of placentas from congenital toxoplasmosis cases in humans is uncommon and no studies that we are aware of quantify trophoblast cell populations at sites of infection. However, the most lethal outcomes of congenital toxoplasmosis occur when infection is acquired by the mother in the first and second trimesters when the placenta is most actively developing. This might suggest a connection between placental age, infection onset, and severity of disease. Moreover, our demonstration of fusion arrest in *Toxoplasma*-infected TS-CTB suggests that areas of infection could result in proximal non-renewing STB. Such weakening of the STB layer might contribute to suppressed nutrient and oxygen exchange, leading to local symptoms of placental insufficiency.

*T. gondii* is unique in its ability to infect nearly all host cell types. There are several examples that demonstrate how its manipulation of diverse host cell functions provides direct advantage to the parasite. For instance, *T. gondii* enhances the motility of infected dendritic cells^21^ and disrupts barrier function in endothelial cells^68^, both of which may aid in its dissemination to distant tissues. In applying this principle to the findings of this study, it is provocative to imagine how or why *Toxoplasma* may have evolved the capacity to disrupt cell differentiation in the placenta. In exploring this, we would like to present two connected concepts: (1) that the evolutionary history of *Toxoplasma* has been written primarily in its cycling through mice and cats whose placentas bear low anatomical and physiological conservation with humans, and that therefore, (2) congenital transmission is more likely a byproduct of the uniquely large host cell range and high dissemination capacity of *Toxoplasma,* rather than an evolved favorable route of survival for the parasite

If we accept this to be the case, it is fascinating to imagine how the manipulation of host cell identity by *T. gondii* could have evolved for its utility beyond the placenta in other body systems. *Toxoplasma* may have originally gained these capabilities through its infection of cell types that are more relevant to its pathogenesis, like dendritic cells or fibroblasts. However, the stemness and fate potential of the cells infected is what dictates the ultimate impact of dysregulation. In the case of placental trophoblasts, the impact is quite severe in that *Toxoplasma* prevents fusion and formation of the robustly antimicrobial syncytiotrophoblast. In other cell types the developmental impact of *T. gondii* infection is not necessarily as extreme, or detected as obviously, due to the absence of downstream developmental fates.

Expanding on this concept, our investigation into differentially enriched motifs in open chromatin regions reveals a significant enrichment of the human AP-1 binding motif “TCA_TGA” in *Tg*-infected cells. This supports what has long been observed in *Toxoplasma* infection of a variety of human cells where AP-1 transcription factors are activated and presumably dictate host-cell transcriptional changes^69,19,70^. The importance of this is two-fold. For one, AP-1 transcription factor complexes and the enhanced expression of their resulting gene targets are broadly implicated in many types of cellular differentiation^71,72^. This includes developmental pathways relevant to neurons, leukocytes, and muscle, cell types which have much higher relevance in the typical pathogenesis of *Toxoplasma*. But perhaps more interestingly, the enrichment of this motif presents an opportunity for other transcription factors with the same cognate recognition motif to bind. For example, BACH2 is highly expressed in cell column cytotrophoblasts, is induced by *Toxoplasma* infection, and uses the same “TGA_TCA” binding motif. It is not unreasonable to think that the same mechanisms that increase AP-1 binding site accessibility during *Toxoplasma* infection might also, even unintentionally, prime the chromatin landscape for developmentally relevant TFs (like BACH2) to initiate cellular differentiation. This could have evolved by selection-driven adaptation, but we find it more likely that infection-driven differentiation is a mechanism that is broadly applied across multiple cell types. As a generalist, it seems likely that *T. gondii* has evolved to target only core host gene networks such that they reliably present in the many cell types that the parasite encounters.

In summary, thanks to strong recent advances in tractable placental models, we can now use an *in vitro* system to map the progression of a specific developmental process and precisely place *T. gondii*-infected cells within that trajectory. Developmentally flexible and tunable systems like trophoblast stem cells and trophoblast organoids provide an unprecedented opportunity to evaluate how *Toxoplasma* infection might influence cell differentiation. Similarly, access to primary chorionic villi provides an essential validation tool to relate *in vitro* findings to what occurs in complex tissue. Beyond novel modeling systems, we have demonstrated new approaches to studying host-parasite interactions both generally and specifically in the context of the human placenta. We take advantage of “contaminating” parasite-derived transcripts in single-cell and single-nucleus sequencing to directly identify infected cells within a biological sample, allowing for discrimination between these cells and uninfected bystanders. We present the technique of blunt serial transverse dissection of Caesarian-derived *ex vivo* chorionic villi to drastically improve the infection rate of underlying CTB. Finally, we present a novel approach to assess functional trophoblast fusion through a three-dimensional acute co-culture of TS-CTB and mature TS-STB. These approaches have defined a new biological consequence of congenital *Toxoplasma* infection that has broad implications for our understanding of disease pathology, evolved parasite dissemination strategies, and the capacity for pathogens to subjugate core developmental processes.

### Limitations of the Study

We have demonstrated how *T. gondii* antagonizes STB development by promoting EVT development in placental trophoblasts. This phenomenon was identified using primarily transcriptomic and morphological data with moderate assessment of protein-level changes via Western blot and Luminex. This study would benefit from single-cell proteomic data to assess the conservation between the transcriptomic changes observed and proteomic changes. We also identify EMT and decidual signaling cues as gene networks that promote EVT formation in trophoblast organoids. Further work is needed to identify nature of the interaction between these gene networks and *Toxoplasma* parasites. Moreover, we propose that this phenomenon predicates parasite transmission to the fetus in cases of clinical congenital toxoplasmosis. No *in vitro* model exists that could replicate this condition in the context of human placental anatomy. Meaningful information could be captured using non-human primate or other animal models of congenital *Toxoplasma* infection to assess whether polarization of trophoblast development leads to higher parasite transmission incidence.

## ACKNOWLEDGEMENTS

The authors would like to acknowledge the various core facilities that contributed to the execution of this work including the Health Sciences Sequencing Core at UPMC Childrens, the Single Cell Sequencing Core at UPMC Presbyterian, and the specimen collection team and the women whose placental tissue was collected at UPMC Magee Womens. The authors would also like to thank the individuals of each of their labs that are not authors of this work for their intellectual contribution and discussion. Funding that supported this work includes grants: R01HD1066247 (JPB, CBC), K12HD000849 (CJM), F31AI167594 (LFC), T32GM133353 (LFC).

## AUTHOR CONTRIBUTIONS

Conceptualization, LFC, JPB; Methodology, LFC, LY, JPB; Formal Analysis, LFC, JPB; Investigation LFC, LY, MG, NNW, SMR, RJS; Resources, JPB, CBC, CJM; Data Curation, LFC. Writing – Original Draft, LFC, JPB; Writing – Review & Editing, LFC, JPB, CBC, LY, SMR. Funding Acquisition, JPB, CBC, CJM, LFC.

## SUPPLEMENTARY FIGURE TITLES/LEGENDS

**Figure S1: *T. gondii* inhibits STB fusion**

(A) Western blot of whole cell lysate from mock-infected and *Tg*-infected TS-CTB at 48 hpi. Proteins visualized using hCG polyclonal antibody. Molecular weights of observed bands likely indicates detection of multiple CGB isoforms. Protein quantified using integrated density of observed bands normalized to histone H3 loading control within timepoint. Significance determined one-sample two-tailed t-test. (B) More images of flox dsRed to eGFP TS-STB co-culture with either mock-infected or *Tg*-infected TS-CTB-cre from experiment in main figure. Red and green channels projected singularly in greyscale or combinatorically in color. Quantification of GFP across conditions by integrated density and normalized to total cell area. Measurements collected at conclusion of 48 hour co-culture. Each point represents an average of 4 fields of view at 4x magnification per single biological replicate for a total of three biological replicates per condition. (C) Quantification of experiment 2 of TS-CTB-cre and floxed TS-STB co-culture. (D) Representative IF images visualizing *T. gondii* in Tg-infected TS-CTB-cre co-culture. (D) Viability assay for 3D TS-CTB-cre at 24, 48, and 72 hpi. Quantification indicates proportion of dye excluding (live) TS-CTB-cre at the given time point. Significance determined by two-way repeated measures ANOVA with Fisher’s LSD correction. For all, * p < 0.05, ** p < 0.01, *** p < 0.001, **** p < .0001, ns = not significant.

**Figure S2: Tg-infected TS-CTB have signatures of TS-EVT development but are not impacted by various Tg-knockout strains.**

(A) Full GO analysis output from Figure 2A. (B) Differential expression of select lineage markers and EMT factors from 24-hour mock- or Tg-infected TS-CTB infections. (C) Scratch assay to determine motility of mock-versus Tg-infected TS-CTB. Gap closure data analyzed by one-sample two-tailed t-test. (D) *ITGA2* and *SDC1* transcript expression analysis via qPCR among mock- and Tg-infected TS-CTB using Type II *T. gondii* (strain = ME49). (E and F) *ITGA2* and *SDC1* transcript expression analysis via qPCR using indicated *T. gondii* knockout strains. For all qPCR, points represent three technical replicates of 1 biological replicate each. One-sample two-tailed unpaired t-test performed on linear ΔCT values and results for significance projected onto fold change (* p < 0.05, ** p < 0.01, *** p < 0.001, **** p < .0001).

**Figure S3: Further analysis of scRNA-seq data indicating Tg-induced developmental disruption.**

(A) Representative phase-contrast images of cells used for sequencing at their respective endpoints. (B) Model putative developmental trajectory, UMAP colored by biological sample origin, and UMAP of all cells colored by pseudotime. (C) TgGRA1(+) cells from Tg-infected TS-CTB biological sample (D) Top 10 and bottom 10 increased/decreased transcripts among TgGRA1(+) cells from Tg-infected TS-CTB sample. Rank ordered by log2FC. (E) Output of all cells mapped against *T. gondii* genome, n = 211. (F) All significantly differentially expressed transcripts between cluster 1 and cluster 2 of *T. gondii*-infected cells. (G) TgRONs expression throughout clusters.

**Figure S4: BACH2 chromatin accessibility increases in Tg-infected TS-CTB**

(A) Coverage plot for BACH2 locus chromatin accessibility and expression between TgGRA1(+) and TgGRA(-) cells in TS-CTB cluster. Linked peaks indicate areas of chromatin accessibility that gain accessibility in a coordinated fashion between samples. (B) Motif footprint plot for BACH2 of TgGRA1(+) cells versus TgGRA1(-) cells in TS-CTB cluster. Tn5 insertion enrichment indicates average accessibility prior to and immediately following motif occurrence.

**Figure S5: GSEA output of Tg-infected versus mock-infected CV**

(A) Left: full GSEA output of Tg-infected versus mock-infected chorionic villi. Right = select enrichment plots for highly ranked hallmark pathways

**Figure S6: *Tg*-infected trophoblast organoids have unique transcriptomic signatures.**

(A) CTB-out orientation of TO1. CK19 (cyan) indicate mononuclear trophoblasts while SDC1 (red) indicates fused STB core. (B) Mock- and Tg-infected TO1 in liquid suspension with increased HLA-G staining but no detectable morphological changes (C) qPCR quantification of *ITGA2* and *SDC1* of mock-infected versus *Tg-*infected TOs. Housekeeping transcript is *GAPDH*. Points represent three averaged technical replicates of one biological replicate. Significance determined through analysis of linear ΔCT values between treatments by one-sample two-tailed t-test (** p < 0.01). Results are projected onto fold change measurements. Data is representative of two independent experiments. (D) (D) PCA of bulk RNA-seq data from TO1-EVTs, TO2 mock/*Tg-*infected, and TO1 mock/*Tg-*infected. Subset PCAs of TO1 mock/*Tg-*infected and TO2 mock/*Tg-*infected. (E) Left: volcano plots of TO2 mock-/Tg-infected and reproduced plot of TS-CTB Tg-infected versus mock. Right: GSEA enrichment plots for Hallmark epithelial to mesenchymal transition. (F) ChEA3 transcription factor enrichment among infection-induced transcripts in TO1.

**Figure S7: *Tg*-infected TOs have a pro-inflammatory secretome that resembles elements of decidual signaling**

(A) GSEA output of *Tg*-infected TO1 versus mock and TO1-EVT versus mock. For all, p < 0.05 (B) Heatmap of 22 analytes from custom Luminex analysis of mock-infected and *Tg-*infected TS-CTB. Data represent log2fold change from DMEM (unconditioned media). (C) Expression of top 15 differentially expressed secreted proteins from DOs in Yang et. al *eLife* between Tg- and mock-infected TO1. Samples with a value of 0 not visualized due to log scale. Data analyzed using multiple unpaired t-tests with Holm-Šídák correction for multiple comparisons. (D) Differential expression of genes involved in decidual signaling that are increased during Tg-infection in at least 2/5 datasets. Table includes respective fold change and p-value significance of each across 5 bulk or snRNA-seq datasets. Fold changes for snRNA-seq are from TgGRA1(+) cells from cluster 1 versus all cells in cluster 1 (putative CTB). For bulk datasets, fold changes and p-values are from Tg-infected samples versus mock-infected samples. * p < 0.05, ** p < 0.01, *** p < 0.001, **** p < 0.0001. (E) Full list of analytes from Figure 7B.

## STAR METHODS

### Lead contact

Further information and requests for resources and reagents should be directed to and will be fulfilled by the lead contact, Leah Cabo.

### Materials availability

Materials generated in this study are available upon reasonable request.

### Data and code availability

All the data generated in this study have been deposted at GEO: Gene Expression Omnibus and are publicly available as of the date of publication. Accession numbers are listed in the key resources table. This paper also analyzes existing, publicly available data. The accession numbers for these datasets are also listed in the key resources table. Codes used for all the analyses are available at: *pending*

### Continuous culture of human trophoblast stem cells

TS cells (clone 27) derived from first trimester placental tissue were generously provided by Dr. Hiroaki Okae of Tohoku University Graduate School of Medicine. Cells are grown as described previously^29^ with some modifications that reflect the adaptation of established protocols to our laboratory environment. In summary, 6-well dishes are seeded with 2 mL of TS cell suspension at a concentration of between 3 x 10^4^ to 4.5 x 10^4^ cells per mL depending on the target density and growth timeline. Cells are suspended in CTB medium consisting of TS basal medium: DMEM/F12 (Gibco, Waltham, MA, USA), 1% ITS-X100 (ThermoFisher Scientific, Waltham, MA, USA), 0.3% fatty acid free bovine serum albumin (BSA) (Sigma, St. Louis, MO, USA), 200 μM ascorbic acid (Sigma), 0.5% penicillin-streptomycin (ThermoFisher Scientific), and 0.5% Knockout Serum Replacement (KSR) (Gibco) supplemented with 25 ng/mL epidermal growth factor (EGF) (ThermoFisher Scientific), 2 μM CHIR99021 (Stemolecule Reprocell USA, Inc., Beltsville, MD, USA), 5 μM A83-01 (Stemolecule Reprocell USA, Inc., Beltsville, MD, USA), 0.8 mM VPA (APExBIO, Houston, TX, USA), 5 μM Y27632 (Stemolecule Reprocell USA, Inc., Beltsville, MD, USA), and 2 μg/mL iMatrix-55 (AMSBIO, Abingdon, UK). Cells are grown at 37°C with 5% CO_2_ until 70-90% confluent before subsequent passage.

To detect broad changes with greater sensitivity and in a more developmentally flexible environment, all experiments (besides the previously published bulk RNA-seq data from Figure 2) were conducted using a media condition consisting of the basal media recipe from Okae *et. Al* plus the ROCK inhibitor Y27632 (unless otherwise noted, for example, for TS-EVT and TS-STB differentiation). These media components are described as the minimum nutritional requirements to continuously culture TS-CTB and allow us to more clearly observe infection-induced changes in differentiation as the cells are not being treated with components that forcibly maintain their stem-cell status. As described in Okae *et. al* and confirmed in our hands, this condition elicits some predisposition towards fusion and a STB-like identity.

### Differentiation of TS-CTB to 3-dimensional TS-STB and 2-dimensional TS-EVT

For 3D TS-STB induction, approximately 1.2 x 10^5^ to 1.5 x 10^5^ cells (< passage 40) per well suspended in 3 mL of TS-STB media (TS differentiation basal: (DMEM/F12 [Gibco], ITS-X100 [ThermoFisher Scientific], 0.3% fatty acid-free BSA [Sigma], 0.5% penicillin-streptomycin [ThermoFisher Scientifc], 0.1 mM 2-mercaptoethanol [Fisher Scientific]), supplemented with 2.5 μM Y27632 (Stemolecule Reprocell), 2 μM forskolin (Sigma), 4% EGF (ThermoFisher) and 50 ng/mL KSR (Gibco)) are seeded in each well of a 6-well dish. Cells should remain non-adherent and will aggregate in the center of the well over time. At day 3 of growth, supplement each well with an additional 2 mL media. TS-STB at either day 5 or day 6 of differentiation were used for experiments.

For 2D TS-EVT induction, 6-well dishes are pre-coated in 1 ug/mL collagen IV. Approximately 7 x 10^5^ cells ( < passage 40) per well are seeded in 2 mL TS-EVT media (TS differentiation basal: (DMEM/F12 [Gibco], ITS-X100 [ThermoFisher Scientific], 0.3% fatty acid-free BSA [Sigma], 0.5% penicillin-streptomycin [ThermoFisher Scientifc], 0.1 mM 2-mercaptoethanol [Fisher Scientific]), supplemented with 100 ng/mL of neuregulin 1 (NRG1), 7.5 μM A83-01, and 2.5 μM Y27632, 4% KSR, and 2% Matrigel (Corning, Corning NY, USA) to make EVT media. On day 3 of EVT differentiation, the media was replaced with EVT media excluding NRG1 and with a reduced Matrigel concentration of 0.5%. At day 6 of growth, the medium was replaced with the EVT media excluding NRG1 and KSR plus a reduced Matrigel concentration of 0.5%. TS-EVT at either day 7 or 8 of differentiation were used for experiments.

### Continuous culture of *Toxoplasma gondii* parasites

*T. gondii* parasites are cultured in human foreskin fibroblasts (HFF) in Dulbecco’s modified Eagle medium (DMEM, ThermoFisher Scientific) plus 100 U/L penicillin/streptomycin, 2 mM L-glutamine, 10% FBS (fetal bovine serum), and 3.7 g NaH_2_CO_3_/L. Cells with intracellular parasites are incubated in 5% CO_2_ at 37°C. To prepare parasites for infection assays, infected HFF monolayers were scraped and syringe-lysed (with 25G and 27G needles) to release tachyzoites. Parasites were filtered through a 5.0 μm syringe filter to remove host cell debris and then pelleted at 800 x g for 10 minutes. Parasites were quantified and MOI was calculated based on the initial number of seeded TS cells (example: for 70,000 TS-CTB seeded, MOI 3 = 210,000 parasites). For mock-infections, the same parasite suspension used for infection was passed through a 0.22 μm syringe filter to remove all parasites. The resulting elutate was used to treat host cells at the same dilution as the parasite suspension.

### Trophoblast organoid culture

Trophoblast organoids were maintained in culture as described in Yang et. al, 2023^57^. TOs at day 5 post passage were used for infection. TOs at day 8 of EVT induction were used for comparison in all presented figures. TO1, TO2, and TO90 represent independently isolated and maintained lines of trophoblast organoids from different patient placentas. TO1 and TO90 were isolated from biologically male fetuses/placentas (XY) whereas TO2 is from biologically female fetuses/placentas (XX). For culture of TOs that would eventually be used for *Tg-*infection, mock-infection and EVT differentiation, TOs were seeded in an 8-well chamber slide on a thin layer of Matrigel (Corning).

### Gene set enrichment analysis (GSEA)

GSEA (MSigDB, UC San Diego and Broad Institute) software was downloaded from https://www.gsea-msigdb.org/gsea/index.jsp and used as indicated in the accompanying documentation. Briefly, log-normalized count data from bulk RNA-seq datasets was extracted using DESeq2 and formatted according to the requirements of GSEA software (expression dataset file format = .gct; phenotype labels file format = .cls). Genes with an expression of 0 across all samples were removed as to not create artificial distance between highly and lowly expressed genes. For all, the gene set collection used was “Human Hallmark gene sets”. Other available human gene set collections were queried for investigation but are not presented in this manuscript. For each run, 1000 permutations were run with the “Human_Gene_Symbol_with_Remapping_MSigDB.v202X.2.Hs.chip” (with X referring to the year in which each analysis was run [2020-2023]). Enrichment statistic used was “weighted” and the metric used for ranking genes was “Signal2Noise”. The max gene set size was 500 and the minimum gene set size was 15. Respective FDR and p-values are reported as calculated by GSEA software.

### Gene ontology analysis

Gene ontology analysis was performed using the web-based tool GO Enrichment Analysis powered by PANTHER (https://geneontology.org). Number of genes queried varies between analysis but generally represents differentially expressed transcripts between conditions with a p-value < 0.05 plus −2 > Log2FC > 2. Resulting enriched GO terms were sorted by lowest FDR and ranked accordingly.

### ChEA3 transcription factor enrichment analysis

The ChEA3 ChIP-X Enrichment Analysis (Version 3) webtool^46^ was used to conduct transcription factor enrichment analysis. Genes queried varies between analysis but generally represents differentially expressed transcripts between conditions with a p-value < 0.05 plus −2 > Log2FC > 2. Results were sorted by lowest Mean Rank across the included datasets (sourced from ENCODE, ReMap, GTEx, ARCHS4, and literature).

### shRNA construct design and ligation to plasmid backbone

Constructs were designed for direct insertion into the pLVTHM plasmid which was a generous gift from Didier Trono^73^ (Addgene plasmid #12247). The TRC (The RNAi Consortium, Broad Institute) hairpin design webtool was used to generate shRNA sense and antisense sequences for a chosen transcript: https://portals.broadinstitute.org/gpp/public/seq/search. Designs were crosschecked using the siRNA Scales webtool: http://gesteland.genetics.utah.edu/siRNA_scales/index.html and the Whitehead Institute siRNA selection program: https://sirna.wi.mit.edu/siRNA_search.cgi?tasto=102653334. For each given construct design, the target GC percentage ranged from 30-52% with no runs of 4 or more A/T/G nucleotides, high thermodynamic value, high predicted knockdown efficiency (>90%), and low chance of off-target effects (< 16 of 19 bp overlapping with non-target mRNA sequence) as determined by BLAST analysis. The loop sequence used for all was CTCGAG with a poly T on the forward oligo and poly A on the reverse oligo. Ends of constructs were designed to re-construct the MluI and ClaI sites used for cloning. pLVTHM was linearized using MluI and ClaI and gel purified. Forward and reverse oligos were ordered in pairs from IDT, annealed, and ligated into cut pLVTHM using standard T4 ligation (NEB). For validation of successful ligation into pLVTHM, we use the new XhoI site introduced in the loop sequence of all shRNA constructs to do a diagnostic restriction enzyme digest with XhoI and PvuI. We also submitted plasmids for Sanger sequencing using the H1 primer from Genewiz (Azenta) to assure successful insertion. For each target transcript, 5 shRNA constructs were designed, cloned, and purified. After individual purification, the 5 hairpin constructs were pooled at an equal concentration for use in lentiviral transfection. This approach results in a mixed population of lentivirus and, therefore, compounded transduction of multiple shRNA constructs into a target cell. Transcript knockdown was validated using qPCR against the target sequence.

### Lentivirus production and cell transduction

High efficiency lentivirus production and cell transduction was optimized using procedures outlined in Pirona et. al 2020^74^. HEK-293T cells were cultured in cDMEM growth medium. Cells were passaged routinely and at a high density between passages. 6-well dishes were seeded to reach 90% confluence prior to transfection. To transfect HEK-293T cells, a DNA solution was made by mixing 1.47 μg purified psPAX2 plasmid (lentiviral packaging plasmid, Addgene), 0.80 μg pMD2.G (VSV-G envelope expressing plasmid, Addgene) and 1.72 μg cargo plasmid (plasmid variable depending on KO or KD approach and target transcript) per well to transfect. The DNA solution was combined with 5 μL Lipofectamine PLUS reagent and 6 μL Lipofectamine P3000 reagent in 250 μL OptiMEM (solution A). A separate solution of 250 μL OptiMEM, 7 μL Lipofectamine 3000 reagent, and 5 μL Lipofectamine LTX reagent was made (solution B). Solutions A and B (per cargo plasmid) were combined and incubated at room temperature for 30 minutes. The combined solution was added dropwise to a well of HEK293-T cells and mixed gently. After 24-48 hours, cell supernatant containing synthesized lentivirus was collected, centrifuged at 500 x g for 10 minutes to exclude non-adherent HEK cells, aliquoted to single use volumes, and frozen at −80°C (or used fresh). To transduce, supernatant containing lentivirus was added, either fresh or having been thawed on ice, to 30-50% confluent target cells. Transduction was allowed to progress for 2-3 days prior to cell passage or drug selection (for certain cargo plasmids). Transduction progress was monitored through fluorescent reporter expression.

### Fusion co-culture assay

Two transgenic TS-CTB cell lines were created using lentiviral constructs deposited at Addgene: pLV-EF1-Cre-PGK-Puro was a generous gift from Javier Alcudia (Addgene plasmid # 108543) and pLV-CMV-LoxP-DsRed-LoxP-eGFP was a generous gift from Jacco van Rheenen (Addgene plasmid # 65726). Lentivirus was produced as described above and TS-CTB were independently transduced with 500 μL of virus-containing supernatant (per construct) prior to 3 days of selection with 2 μg/mL puromycin. Cells were removed from selection and cultured in normal conditions for a maximum of 5 passages. The resulting constitutively red-fluorescing “TS-CTB-flox” [TS-CTB-CMV-LoxP-DsRed-LoxP-eGFP] were differentiated to 3D TS-STB as described. Simultaneously, the resulting non-fluorescent “TS-CTB-cre” [TS-CTB-EF1-Cre-PGK-Puro] were infected with MOI 3 (210,000 parasites) WT RH88 (nonfluorescent, WT parasites) or mock-infected in 2-dimensional culture conditions overnight (TS basal + Y27632 medium). *Tg-* infected or mock-infected TS-CTB-cre were trypsinized, centrifuged, and resuspended in new TS-basal + Y27632 medium. Simultaneously, TS-STB-flox were pelleted, resuspended in TS-basal + Y27632 medium, and redeposited in wells of a new 6-well dish. Either *Tg-*infected or mock-infected TS-CTB-cre were added to TS-STB-flox in acute co-culture. All cells were co-cultured for 48 hours and fluorescent images were taken while cells were in an unfixed, live state. Fusion(+) TS-STB-flox + TS-CTB-cre hybrid cells were determined by the co-fluorescence of red and green while fusion(-) TS-STB-flox cells were determined by the sole fluorescence of red. Non-fluorescent cells in images are unfused *Tg-*infected or mock-infected TS-CTB-cre.

### Viability assay

To test the viability of TS-CTB in 3-dimensional non-adherant culture with and without *T. gondii* infection in the same conditions of the 3D fusion assay, stable TS-CTB-cre were infected with MOI 3 (70,000 cells plated = 210,000 parasite) RH/YFP parasites for 24 hours in TS basal + Y27632 media in 6-well dishes. 100 μL of cell suspension was removed from each well and combined. Cell viability was assessed using 0.4% Trypan Blue Solution (ThermoFisher) and a manual hemocytometer. Cells staining blue were considered dead whereas phase-bright non-blue cells were considered alive. *Tg-*infected and mock-infected TS-CTB-cre were then removed from the plate using 1:1 TrypLE Express (ThermoFisher) and 1X PBS. Cells were centrifuged at 250 x g for 5 minutes, washed with PBS, and resuspended in new TS basal + Y27632 media and deposited in a new 6-well dish. At 48 hours post infection, cell suspension was again sampled as before using 200 μL from each well between treatments. At 72 hours post infection, all cells were removed from each well, centrifuged, and resuspended for counting. Number of cells counted at 72 hpi represents a fraction of total cells.

### Single cell RNA-sequencing using combinatorial barcoding and data analysis

Single cell RNA-sequencing of TS cells was conducted using Parse Biosciences (at time of ordering, SplitBio) split-pool combinatorial barcoding technology. This approach allowed us to retain large, multinucleated 3D TS-STB and fix + store passage-matched endpoint-distinct samples. First, we optimized cell fixation and sample preparation to maximize the quality and quantity of input cell samples. Cells were cultured and infected as described above. Mock-infected and *Tg-*infected TS-CTB were infected 24 hours after passage for 48 more hours. Cells were fixed and stored according to Parse protocols. Passage-matched TS-STB and TS-EVT were differentiated for 6 and 8 days respectively (as described) and fixed and stored according to Parse protocols. Next, we completed the split-pool combinatorial barcoding and library prep pipeline as indicated by the manufacturer’s protocols with the important exception that we did not use cell filters at any point as to not exclude the comparably larger TS-STB. We sequenced 1 of the 8 sublibraries produced by the kit to yield our 10,622 cell dataset with 200 million reads (50M per sample, combined into one library). Data was analyzed using Parse Biosciences “split-pipe” Linux-based pipeline to deconvolute individual cells and map against a custom genome. The *Toxoplasma GRA1* gene (“TgGRA1”) was manually added to the hg38 human genome to produce a hybridized genome. The resulting filtered gene matrix was imported and analyzed using Seurat in R. Strict cell filtering cutoffs were used: 100 < RNA features < 3500, 1000 < RNA counts < 15000, % mitochondrial reads < 25. TgGRA1(+) cells belonging to any biological sample preparation that did not experience infection (mock-infected TS-CTB, TS-STB, and TS-EVT) were also filtered out as putative doublets. Data were normalized and dimensions 1-20 were used for clustering. Code used to analyze the dataset is publicly available at: *pending* and data is available on GEO under accession record: *pending*

### Single nucleus multiomic sequencing and data analysis

Passage-matched mock-infected and *Tg*-infected TS-CTB, Day 4 TS-EVT, and Day 8 TS-EVT samples were generated as described. Nuclear extraction was performed according to 10x Genomics nuclear isolation protocol for multiome sequencing. Samples were sequenced with a target of 10,000 cells per sample. Data was aggregated and mapped using 10x Genomics aggr and cellranger pipelines and exported for use in R with Signac (snATAC-seq) and Seruat (snRNA-seq). Complete code used for analysis is available at *pending* and data is available on GEO under accession record: *pending*

### Chorionic villi acquisition, preparation, infection, and RNA extraction

Fresh placental tissues from third trimester caesarian section procedures were obtained from the Steve N. Caritis Magee Obstetric Maternal and Infant Biobank who functioned as an honest broker according to University of Pittsburgh IRB protocol #1900322. Inclusion criteria include patients without labor, or ruptured membranes undergoing cesarean delivery. Data presented in this manuscript represents tissues collected from women between 36-40 weeks of gestation in absence of diagnosed placental pathology or placental disease. Some samples were derived from women with diet-controlled gestational diabetes, and these cases are noted in the text. Tissue samples were obtained within 30 minutes of delivery and were kept submerged in warm DMEM during processing. Chorionic villi were removed from visible decidua and were dissected with aid of a light microscope into wells of a 24-well tissue culture dish. Villi were allowed to rest for >4 hours in media at 37°C with 5% CO_2_ before being resuspended in TSC basal + Y27632 media and serially transversely sliced with dissection scissors to disrupt the syncytiotrophoblast layer and reduce the structure of villi. Villi were infected or mock-infected with 10 million RH/YFP parasites as described above. Infections progressed for 48 hours at which time villi were collected and stored in RNAlater (ThermoFisher) at −80°C. For RNA isolation, villi were thawed on ice and resuspended in RLT buffer (Qiagen). Tissue was homogenized in RLT buffer with the TissueRupter II (Qiagen), a handheld rotor-stator homogenizer. Resulting homogenate was used for total RNA extraction using Qiashredder and RNeasy columns (Qiagen) according to the manufacturer’s instructions.

### Quantitative PCR (qPCR)

To quantify transcript changes within host cells, whole cell RNA lysates were isolated from infected and mock-infected samples using RLT buffer (Qiagen) supplemented with 2-mercaptoethanol. Total RNA was extracted using both QIAshredder and RNeasy columns (Qiagen) and cDNA was produced from 500 ng total RNA using iScript cDNA synthesis kit (Bio-Rad) all following the manufacturer’s instructions. RNA was analyzed via gel electrophoresis and nanodrop quantification. Primers against target transcripts, in addition to human GAPDH which was used as the housekeeping gene, can be found in supplementary methods. All reactions were completed in technical triplicate using a QuantStudio 3 Real-Time PCR System (ThermoFisher). cDNA was diluted 1:10 with water after synthesis. 3 μL diluted cDNA was mixed with 7 μL mastermix [5 μL SYBER green, 0.5 μL 10 μM F primer, 0.5 μL 10 μM R primer, 1 μL water] and genes were amplified using a standard protocol (95°C for 10 min and 40 cycles of 95°C for 15s and 60°C for 1 min). Data were analyzed with QuantStudio Design & Analysis Software. For all, multiple (> or equal to 2) experiments were performed in technical triplicate. Delta CT values (query gene CT – housekeeping gene CT; a linear measurement) were used for statistical analysis and then converted to fold-change difference using the 2^ΔΔCT^ method. Results of statistical analysis with ι1CT values is projected onto fold-change measurements for purposes of data visualization.

### Immunofluorescent assay and image quantification

Cells in were fixed using 4% paraformaldehyde (PFA, ThermoFisher Scientific) for 15 minutes, rinsed with sterile PBS, and permeabilized with 0.25% Triton-X in PBS for 20 minutes. Cells were blocked with 5% BSA in PBS for minimum of 1 hour. Cells were then incubated with each respective antibody at the appropriate working concentration for either 1 hour at room temperature or overnight at 4°C with gentle shaking. Antibodies used in this study can be found in detail below. After primary antibody incubation, cells were washed extensively with PBS, then incubated with appropriate secondary antibodies at 1:1000 dilution and DAPI at 300 ng/mL dilution respectively in 5% BSA-PBS for 1 hour at room temperature. At conclusion of staining of cells in 6-well dishes, new sterile PBS was deposited on wells used for imaging. All imaging was captured same day for cells in 6-well dishes. The same procedure was used for staining non-adherent cells in suspension (ex: for fusion assay) except cells were centrifuged between steps and pellet was resuspended each time. Cells stained in suspension were mounted by resuspending the final pellet in a small volume (<20 μL) mounting media (Vectashield plus DAPI or homemade [9 parts glycerol to 1 part PBS]) and depositing onto a charged glass slide with coverslip. Most images were taken using an Olympus IX83 epifluorescence microscope and X-Cite LED Boost camera with a 4x, 10x or 20x objective lens. Some images were taken using a Nikon A1R inverted confocal microscope at 20x or 40x with air, or 60x or 100x with oil immersion.

| Antibody (supplier, catalog) | Dilution | Methodology |
| --- | --- | --- |
| SDC1 (Abcam, ab128936) | 1:500 | RT for 1 hour OR 4°C overnight |
| ITGA6 (ThermoFisher MA5-16884) | 1:1000 | RT for 1 hour OR 4°C overnight |
| HLA-G (Abcam, ab52454) | 1:200 | 4°C overnight ONLY |
| ITGA2 (R&D, MAB1233) | 1:1000 | RT for 1 hour OR 4°C overnight |
| Cytokeratin 19 (Abcam, ab52625) | 1:1000 | RT for 1 hour OR 4°C overnight |
| Toxoplasma gondii (Thermo, PA1-7256) | 1:1000 | RT for 1 hour ONLY |
| Alexa Fluor 594, 647, and 488-conjugated host specific secondary aBs (donkeyαgoat, goatαmouse, goatαrat, goatαrabbit) | 1:1000 | RT for 1 hour ONLY |

### Disruption of *T. gondii* candidate genes and mutant strain generation

Specific guide RNAs (gRNAs) were designed using EuPaGDT design tools for both genes. Through Gibson Assembly, the individual gRNAs were inserted into a version of the pSag1::Cas9::GFP::DHFR plasmid that was generously provided by the Treeck lab^75^. This plasmid contained a T2A skip peptide fusing a nuclear localization signal, GFP, Sag1 gene marker, HA tag, and DHFR mutated drug marker into one protein. Successful gRNA insertion was validated using whole plasmid sequencing via Plasmidsaurus. Approximately 10,000,000 parasites were transfected with a 10 ug of NotI-HF linearized plasmid using the BTX electroporation system – Electro Cell Manipulator 600. Transfection reactions were transferred directly onto confluent monolayers of HFFs within T25s and left to incubate for 24 hours. After 24 hours of incubation, the infected host cells were supplemented with 5ul of 10mM pyrimethamine in 50mL of cDMEM (final concentration = 1 µM) to select parasites that had successfully incorporated the plasmid. After populations of parasites grew stably in selection media for 2 weeks, clonal populations were obtained through limiting dilution. Parasites were diluted from 3,000 or 4,000 parasites/mL in a total of 10 mL of selection media. Parasites were then dispersed onto confluent HFF monolayers from the center of the 96-well plate to the outermost column to increase the probability of obtaining a single plaque. Plates were left to sit undisturbed for 8 days, and single plaques were chosen and transferred to confluent HFF monolayers in 24-well plates for 7 days. Parasites were then passed into confluent HFF monolayers of T25 flasks. Genomic DNA was obtained from the remaining parasites using the GeneJET genomic DNA isolation kit to genotype the parasites. Genotyping was done by PCR to verify the disruption of the candidate genes loci through insertion of the Cas9 plasmid into the protospacer adjacent motif (PAM) site. Successful plasmid insertion was visualized on a 1% agarose gel that ran at 100V for 40-50 min, and in some cases were sequenced for gene knockout confirmation.

### SDS-PAGE gel and Western blot

Whole cell lysate was collected from samples using 1X Laemmli buffer (Bio-Rad) supplemented with 2-mercaptoethanol. Samples were boiled using a dry bath at 95°C for 10-15 minutes before loading into wells of a 4-20% TGX precast polyacrylamide gel (Bio-Rad). Loaded gels were run using the Mini-PROTEAN Electrophoresis system (Bio-Rad) at 90V for 1-1.5 hours or until the dye front reached the bottom of the gel. Proteins were transferred from the SDS gel to nitrocellulose membrane at 4°C with stirring using 100V for 1.5 hours. Membranes were subsequently blocked with 5% nonfat dry milk dissolved in PBS-Tween20 for 1 hour at room temperature. To visualize CGB protein, membranes were incubated in CGB antibody (1:2000, Abcam ab243581) diluted in 5% nonfat dry milk dissolved in PBS-Tween20 at room temperature for 1 hour. Secondary antibody used was HRP-conjugated α.rabbit IgG diluted in 5% nonfat dry milk dissolved in PBS-Tween20 for 1 hour at room temperature. Membranes were washed extensively with PBS-Tween20 between incubations. Finally, membranes were incubated for 5 minutes with fresh SuperSignal West Pico PLUS Chemiluminescent substrate (Thermo) and visualized.

### Bulk RNA-seq library preparation, sequencing, and analysis

Total mRNA extracted from various samples was prepared into bulk RNA-sequencing libraries using NEBNext Ultra II Directional RNA Library Prep Kit for Illumina (NEB) according to the manufacturer’s instructions. For all, 150-500 ng of total RNA was used for input and NEBNext Multiplex Oligos for Illumina were used for sequencing primers. All samples were sequenced on a NextSeq (Illumina) with 40-65 million reads per sample. Fastq files were uploaded to CLC Genomics Workbench and mapped against the hg38 human genome. Resulting mapped reads were analyzed in R using DESeq2 package for differential expression analysis. Visualizations made with EnhancedVolcano, ggplot, and other packages. Code used for analysis is available at *pending* and data is available on GEO using accession records: *pending* (chorionic villi) and *pending* (trophoblast organoid).

### ELISA

ELISA against human NRG1 (Thermo) was used according to the manufacturer’s instructions. Samples were derived from supernatants collected from various infected and mock-infected cell types stored at −80°C and thawed on ice. Each sample was diluted 1:2 and results were extrapolated to a standard curve of purified NRG1.

### Scratch assay for cell motility

Parasite (RH/YFP) suspension for infection was prepared as described above. Although variable initial cell number was used in this study, MOI, infection duration, and wound-healing duration remain consistent. A MOI = 4 (for example, for 70,000 cells plated = 280,000 parasites used) was used to infect partially confluent CTB where cells are 80-90% confluent in a 6-well dish. Wells with large, overly dense cell clusters were excluded. Prior to infection, culture media was replaced with TS basal + Y27632 media and used throughout. After 6 hours of infection, 3 scratches were made randomly using a P20 pipette tip (same tip was used for all wells of each group). Floating cells were aspirated and TS basal + Y27632 media was resupplied. Images were taken using an Olympus microscope with an automated stage and software to image each position automatically. Positions were stored and measurements represent the same positions analyzed over time. Gap area was measured in pixels by imageJ with a wound-healing plugin and gap closure rate was calculated using the equation (A_t0_-A_t24_)/ A_t0_ (Suarez-Arnedo et al., 2020^76^). Experiments (> 2) were done in biological triplicate and technical triplicate (3 wells, 3 scratches each) and the TS cell passage number for each experiment is no more than P40.

## REFERENCES

1. Megli, C. J. & Coyne, C. B. Infections at the maternal–fetal interface: an overview of pathogenesis and defence. Nat. Rev. Microbiol. 20, 67–82 (2022).

2. McAuley, J. B. Congenital Toxoplasmosis. J. Pediatr. Infect. Dis. Soc. 3, S30–S35 (2014).

3. Bollani, L. et al. Congenital Toxoplasmosis: The State of the Art. Front. Pediatr. 10, 894573 (2022).

4. Daher, D. et al. Comprehensive Overview of Toxoplasma gondii-Induced and Associated Diseases. Pathogens 10, 1351 (2021).

5. Singh, S. Congenital toxoplasmosis: Clinical features, outcomes, treatment, and prevention. Trop. Parasitol. 6, 113–122 (2016).

6. da Silva, R. J. et al. The trophoblast surface becomes refractory to adhesion by congenitally transmitted Toxoplasma gondii and Listeria monocytogenes during cytotrophoblast to syncytiotrophoblast development. mSphere 0, e00748–23 (2024).

7. Robbins, J. R., Zeldovich, V. B., Poukchanski, A., Boothroyd, J. C. & Bakardjiev, A. I. Tissue Barriers of the Human Placenta to Infection with Toxoplasma gondii. Infect. Immun. 80, 418–428 (2012).

8. Ander, S. E. et al. Human Placental Syncytiotrophoblasts Restrict Toxoplasma gondii Attachment and Replication and Respond to Infection by Producing Immunomodulatory Chemokines. mBio 9, e01678–17 (2018).

9. Robbins, J. R., Skrzypczynska, K. M., Zeldovich, V. B., Kapidzic, M. & Bakardjiev, A. I. Placental Syncytiotrophoblast Constitutes a Major Barrier to Vertical Transmission of Listeria monocytogenes. PLOS Pathog. 6, e1000732 (2010).

10. Johnson, L. J. et al. Human Placental Trophoblasts Infected by Listeria monocytogenes Undergo a Pro-Inflammatory Switch Associated With Poor Pregnancy Outcomes. Front. Immunol. 12, (2021).

11. Rollman, T. B. et al. Human trophoblast stem cells restrict human cytomegalovirus replication. J. Virol. 0, e01935–23 (2024).

12. Koi, H. et al. Syncytiotrophoblast Is a Barrier to Maternal-Fetal Transmission of Herpes Simplex Virus1. Biol. Reprod. 67, 1572–1579 (2002).

13. Krop, J., Tian, X., van der Hoorn, M.-L. & Eikmans, M. The Mac Is Back: The Role of Macrophages in Human Healthy and Complicated Pregnancies. Int. J. Mol. Sci. 24, 5300 (2023).

14. Pollheimer, J., Vondra, S., Baltayeva, J., Beristain, A. G. & Knöfler, M. Regulation of Placental Extravillous Trophoblasts by the Maternal Uterine Environment. Front. Immunol. 9, (2018).

15. Arutyunyan, A. et al. Spatial multiomics map of trophoblast development in early pregnancy. Nature 616, 143–151 (2023).

16. Li, Q. et al. Human uterine natural killer cells regulate differentiation of extravillous trophoblast early in pregnancy. Cell Stem Cell 31, 181–195.e9 (2024).

17. Lee, C. Q. E. et al. Integrin α2 marks a niche of trophoblast progenitor cells in first trimester human placenta. Dev. Camb. Engl. 145, dev162305 (2018).

18. Faral-Tello, P., Pagotto, R., Bollati-Fogolín, M. & Francia, M. E. Modeling the human placental barrier to understand Toxoplasma gondiís vertical transmission. Front. Cell. Infect. Microbiol. 13, 1130901 (2023).

19. Blader, I. J., Manger, I. D. & Boothroyd, J. C. Microarray Analysis Reveals Previously Unknown Changes in *Toxoplasma gondii*-infected Human Cells*. J. Biol. Chem. 276, 24223–24231 (2001).

20. Kim, S.-K., Fouts, A. E. & Boothroyd, J. C. Toxoplasma gondii Dysregulates IFN-γ-Inducible Gene Expression in Human Fibroblasts: Insights from a Genome-Wide Transcriptional Profiling1. J. Immunol. 178, 5154–5165 (2007).

21. Sangaré, L. O. et al. In Vivo CRISPR Screen Identifies TgWIP as a Toxoplasma Modulator of Dendritic Cell Migration. Cell Host Microbe 26, 478–492.e8 (2019).

22. Braun, L. et al. A Toxoplasma dense granule protein, GRA24, modulates the early immune response to infection by promoting a direct and sustained host p38 MAPK activation. J. Exp. Med. 210, 2071–2086 (2013).

23. Panas, M. W. & Boothroyd, J. C. Toxoplasma Uses GRA16 To Upregulate Host c-Myc. mSphere 5, 10.1128/msphere.00402-20 (2020).

24. Feldman, T. P., Ryan, Y. & Egan, E. S. Plasmodium falciparum infection of human erythroblasts induces transcriptional changes associated with dyserythropoiesis. Blood Adv. 7, 5496–5509 (2023).

25. Neveu, G. et al. Plasmodium falciparum sexual parasites develop in human erythroblasts and affect erythropoiesis. Blood 136, 1381–1393 (2020).

26. Liempi, A., et al. *Trypanosoma cruzi* induces trophoblast differentiation: A potential local antiparasitic mechanism of the human placenta? Placenta 35, 1035– 1042 (2014).

27. Medina, L. et al. Ex Vivo Infection of Human Placental Explants by Trypanosoma cruzi Reveals a microRNA Profile Similar to That Seen in Trophoblast Differentiation. Pathogens 11, 361 (2022).

28. Varberg, K. M., et al. ASCL2 reciprocally controls key trophoblast lineage decisions during hemochorial placenta development. Proc. Natl. Acad. Sci. 118, e2016517118 (2021).

29. Okae, H. et al. Derivation of Human Trophoblast Stem Cells. Cell Stem Cell 22, 50–63.e6 (2018).

30. Karvas, R. M. et al. Stem-cell-derived trophoblast organoids model human placental development and susceptibility to emerging pathogens. Cell Stem Cell 29, 810–825.e8 (2022).

31. Dong, C. et al. A genome-wide CRISPR-Cas9 knockout screen identifies essential and growth-restricting genes in human trophoblast stem cells. Nat. Commun. 13, 2548 (2022).

32. Bayer, A. et al. Type III Interferons Produced by Human Placental Trophoblasts Confer Protection against Zika Virus Infection. Cell Host Microbe 19, 705–712 (2016).

33. Corry, J., Arora, N., Good, C. A., Sadovsky, Y. & Coyne, C. B. Organotypic models of type III interferon-mediated protection from Zika virus infections at the maternal–fetal interface. Proc. Natl. Acad. Sci. 114, 9433–9438 (2017).

34. Wu, H. et al. Zika virus targets human trophoblast stem cells and prevents syncytialization in placental trophoblast organoids. Nat. Commun. 14, 5541 (2023).

35. Chen, J. et al. A placental model of SARS-CoV-2 infection reveals ACE2-dependent susceptibility and differentiation impairment in syncytiotrophoblasts. Nat. Cell Biol. 25, 1223–1234 (2023).

36. Białas, P., Śliwa, A., Szczerba, A. & Jankowska, A. The Study of the Expression of CGB1 and CGB2 in Human Cancer Tissues. Genes 11, 1082 (2020).

37. Fiddes, J. C. & Goodman, H. M. The gene encoding the common alpha subunit of the four human glycoprotein hormones. J. Mol. Appl. Genet. 1, 3–18 (1981).

38. Mi, S. et al. Syncytin is a captive retroviral envelope protein involved in human placental morphogenesis. Nature 403, 785–789 (2000).

39. Vargas, A. et al. Syncytin-2 plays an important role in the fusion of human trophoblast cells. J. Mol. Biol. 392, 301–318 (2009).

40. Li, X., Li, Z.-H., Wang, Y.-X. & Liu, T.-H. A comprehensive review of human trophoblast fusion models: recent developments and challenges. Cell Death Discov. 9, 1–14 (2023).

41. Subramanian, A. et al. Gene set enrichment analysis: A knowledge-based approach for interpreting genome-wide expression profiles. Proc. Natl. Acad. Sci. 102, 15545–15550 (2005).

42. Ackerman, W. E. et al. Epigenetic changes regulating the epithelial-mesenchymal transition in human trophoblast differentiation. 2024.07.02.601748 Preprint at 10.1101/2024.07.02.601748 (2024).

43. Franco, M. et al. A Novel Secreted Protein, MYR1, Is Central to Toxoplasma’s Manipulation of Host Cells. mBio 7, 10.1128/mbio.02231-15 (2016).

44. Cygan, A. M. et al. Coimmunoprecipitation with MYR1 Identifies Three Additional Proteins within the Toxoplasma gondii Parasitophorous Vacuole Required for Translocation of Dense Granule Effectors into Host Cells. mSphere 5, e00858–19 (2020).

45. Camejo, A. et al. Identification of three novel Toxoplasma gondii rhoptry proteins. Int. J. Parasitol. 44, 147–160 (2014).

46. Keenan, A. B. et al. ChEA3: transcription factor enrichment analysis by orthogonal omics integration. Nucleic Acids Res. 47, W212–W224 (2019).

47. Varberg, K. M. et al. Extravillous trophoblast cell lineage development is associated with active remodeling of the chromatin landscape. Nat. Commun. 14, 4826 (2023).

48. Liu, L. et al. Smad2 and Smad3 have differential sensitivity in relaying TGFβ signaling and inversely regulate early lineage specification. Sci. Rep. 6, 21602 (2016).

49. Yao, C. et al. BACH2 enforces the transcriptional and epigenetic programs of stem-like CD8+ T cells. Nat. Immunol. 22, 370–380 (2021).

50. Jeyarajah, M. J., et al. The multifaceted role of GCM1 during trophoblast differentiation in the human placenta. Proc. Natl. Acad. Sci. 119, e2203071119 (2022).

51. Miller, S. A. et al. LSD1 and Aberrant DNA Methylation Mediate Persistence of Enteroendocrine Progenitors That Support BRAF-Mutant Colorectal Cancer. Cancer Res. 81, 3791–3805 (2021).

52. Rudzki, E. N. et al. Toxoplasma gondii GRA28 Is Required for Placenta-Specific Induction of the Regulatory Chemokine CCL22 in Human and Mouse. mBio 12, e01591–21 (2021).

53. Zhao, H.-J. et al. Bone morphogenetic protein 2 promotes human trophoblast cell invasion by upregulating N-cadherin via non-canonical SMAD2/3 signaling. Cell Death Dis. 9, 174 (2018).

54. You, J. et al. The BMP2 Signaling Axis Promotes Invasive Differentiation of Human Trophoblasts. Front. Cell Dev. Biol. 9, 607332 (2021).

55. Sharma, S., Godbole, G. & Modi, D. Decidual Control of Trophoblast Invasion. Am. J. Reprod. Immunol. N. Y. N 1989 75, 341–350 (2016).

56. Garcia-Alonso, L. et al. Mapping the temporal and spatial dynamics of the human endometrium in vivo and in vitro. Nat. Genet. 53, 1698–1711 (2021).

57. Yang, L. et al. Innate immune signaling in trophoblast and decidua organoids defines differential antiviral defenses at the maternal-fetal interface. eLife 11, e79794 (2022).

58. Yang, L. et al. Innate immune signaling in trophoblast and decidua organoids defines differential antiviral defenses at the maternal-fetal interface. eLife 11, e79794.

59. Turco, M. Y. et al. Trophoblast organoids as a model for maternal–fetal interactions during human placentation. Nature 564, 263–267 (2018).

60. Haider, S. et al. Self-Renewing Trophoblast Organoids Recapitulate the Developmental Program of the Early Human Placenta. Stem Cell Rep. 11, 537–551 (2018).

61. Wakeland, A. K. et al. Hypoxia Directs Human Extravillous Trophoblast Differentiation in a Hypoxia-Inducible Factor–Dependent Manner. Am. J. Pathol. 187, 767–780 (2017).

62. Bao, H. et al. Hyperactivated Wnt-β-catenin signaling in the absence of sFRP1 and sFRP5 disrupts trophoblast differentiation through repression of Ascl2. BMC Biol. 18, 151 (2020).

63. Ma, Q. et al. Extracellular vesicles secreted by human uterine stromal cells regulate decidualization, angiogenesis, and trophoblast differentiation. Proc. Natl. Acad. Sci. 119, e2200252119 (2022).

64. Menkhorst, E. M. et al. Decidual-Secreted Factors Alter Invasive Trophoblast Membrane and Secreted Proteins Implying a Role for Decidual Cell Regulation of Placentation. PLOS ONE 7, e31418 (2012).

65. Fock, V. et al. Neuregulin-1-mediated ErbB2–ErbB3 signalling protects human trophoblasts against apoptosis to preserve differentiation. J. Cell Sci. 128, 4306–4316 (2015).

66. Schleiss, M. R., Aronow, B. J. & Handwerger, S. Cytomegalovirus Infection of Human Syncytiotrophoblast Cells Strongly Interferes with Expression of Genes Involved in Placental Differentiation and Tissue Integrity. Pediatr. Res. 61, 565–571 (2007).

67. Tabata, T. et al. Human cytomegalovirus infection interferes with the maintenance and differentiation of trophoblast progenitor cells of the human placenta. J. Virol. 89, 5134–5147 (2015).

68. Franklin-Murray, A. L., et al. Toxoplasma gondii Dysregulates Barrier Function and Mechanotransduction Signaling in Human Endothelial Cells. mSphere 5, e00550-19 (2020).

69. Saeij, J. P. J. et al. Toxoplasma co-opts host gene expression by injection of a polymorphic kinase homologue. Nature 445, 324–327 (2007).

70. Gail, M., Gross, U. & Bohne, W. Transcriptional profile of Toxoplasma gondii-infected human fibroblasts as revealed by gene-array hybridization. Mol. Genet. Genomics 265, 905–912 (2001).

71. Katagiri, T., Kameda, H., Nakano, H. & Yamazaki, S. Regulation of T cell differentiation by the AP-1 transcription factor JunB. Immunol. Med. 44, 197–203 (2021).

72. Novoszel, P. et al. The AP-1 transcription factors c-Jun and JunB are essential for CD8α conventional dendritic cell identity. Cell Death Differ. 28, 2404–2420 (2021).

73. Wiznerowicz, M. & Trono, D. Conditional suppression of cellular genes: lentivirus vector-mediated drug-inducible RNA interference. J. Virol. 77, 8957–8961 (2003).

74. Pirona, A. C., Oktriani, R., Boettcher, M. & Hoheisel, J. D. Process for an efficient lentiviral cell transduction. Biol. Methods Protoc. 5, bpaa005 (2020).

75. Young, J. et al. A CRISPR platform for targeted in vivo screens identifies Toxoplasma gondii virulence factors in mice. Nat. Commun. 10, 3963 (2019).

76. Suarez-Arnedo, A. et al. An image J plugin for the high throughput image analysis of in vitro scratch wound healing assays. PLOS ONE 15, e0232565 (2020).

77. Moreau, P. et al. HLA-G gene transcriptional regulation in trophoblasts and blood cells: differential binding of nuclear factors to a regulatory element located 1.1 kb from exon 1. Hum. Immunol. 52, 41–46 (1997).

