## supplemental figures for "*Toxoplasma gondii* infection misdirects placental trophoblast lineage specification"

Supplemental Figure 1
Click here to access/download;Supplemental Figure;figureS1.pdf

### Figure S1

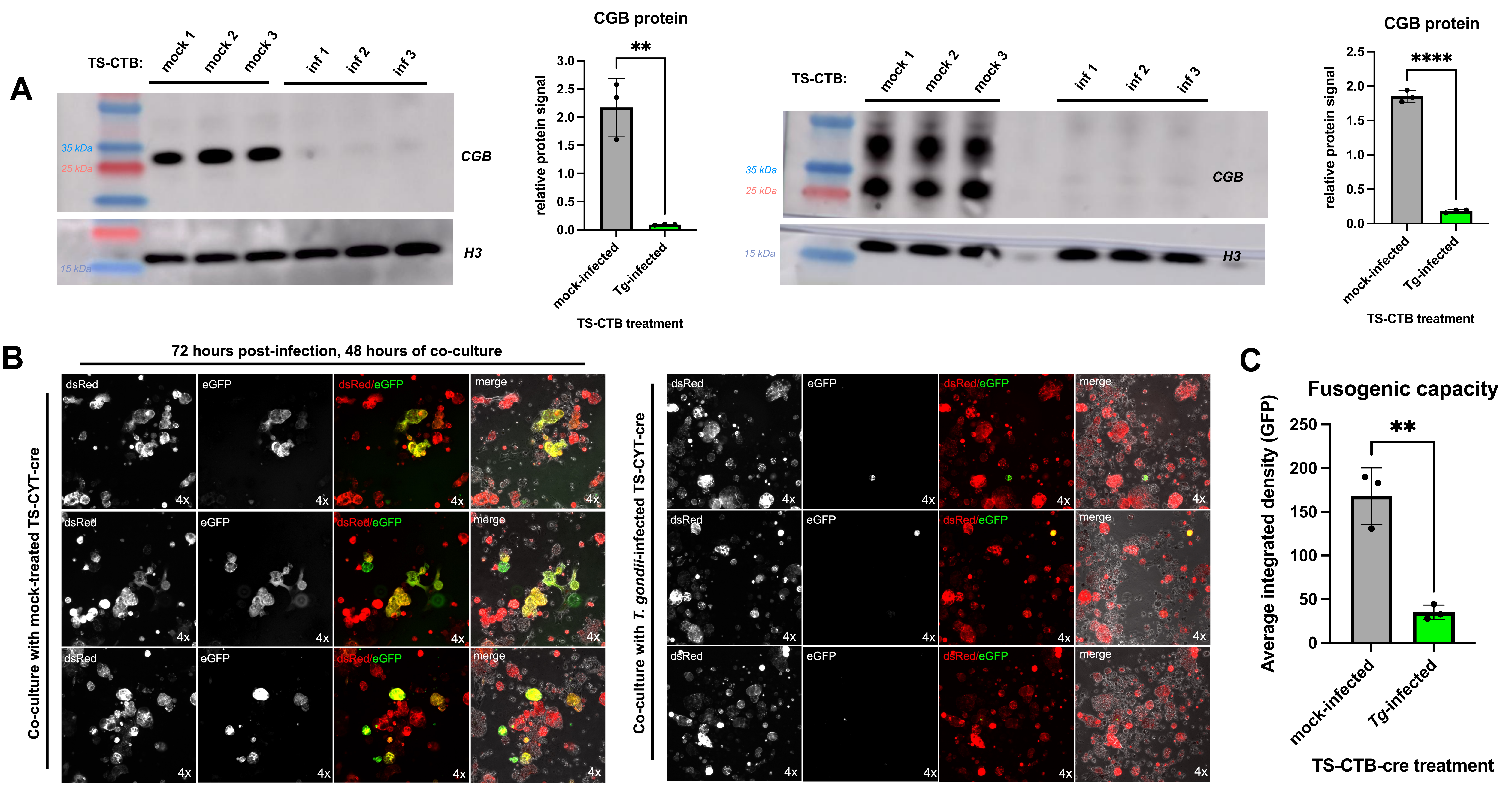


D

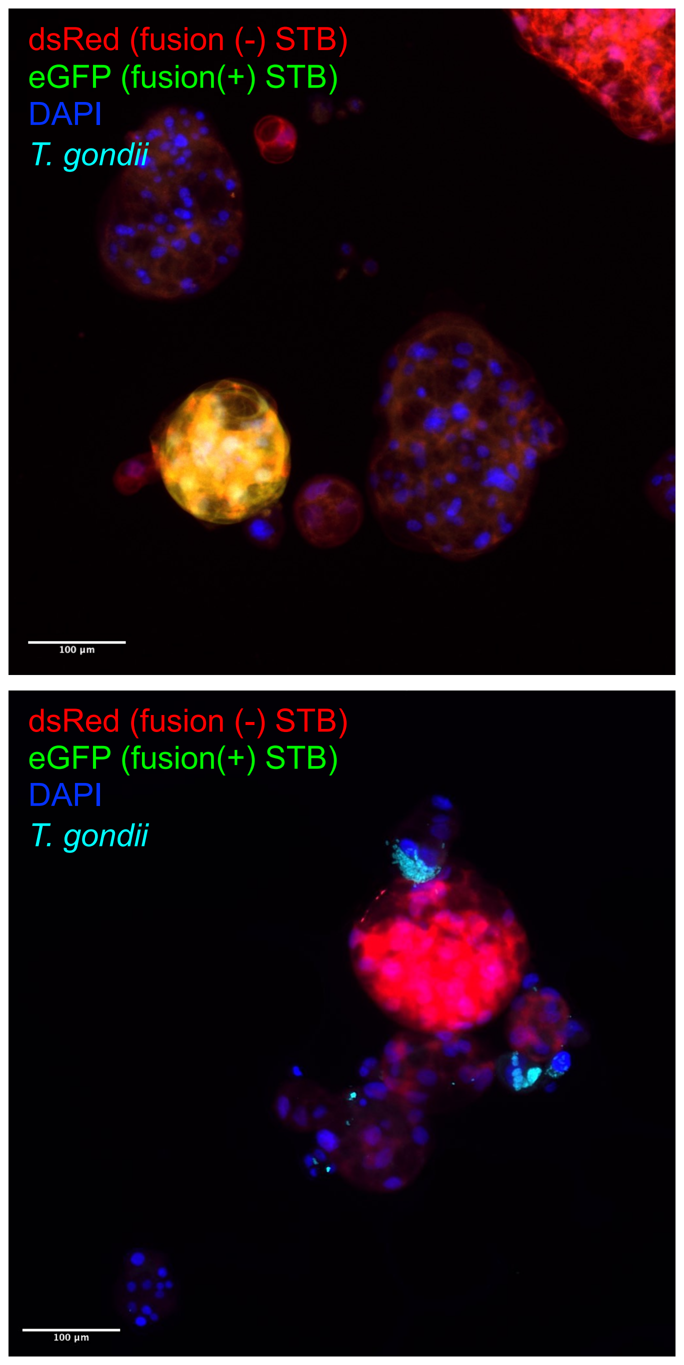


E

| Timepoint (hours post infection) | Condition | Total cells counted | Live cells | Dead cells | % Live | % Dead |
|---|---|---|---|---|---|---|
| 24 hpi | Mock | 15 | 15 | 0 | 100% | 0% |
| | Infected | 31 | 28 | 3 | 90.3% | 9.7% |
| 48 hpi | Mock | 46 | 33 | 13 | 71.7% | 28.3% |
| | Infected | 45 | 35 | 10 | 77.8% | 22.3% |
| 72 hpi | Mock | 662 | 507 | 155 | 76.6% | 23.4% |
| | Infected | 680 | 474 | 206 | 69.7% | 30.3% |

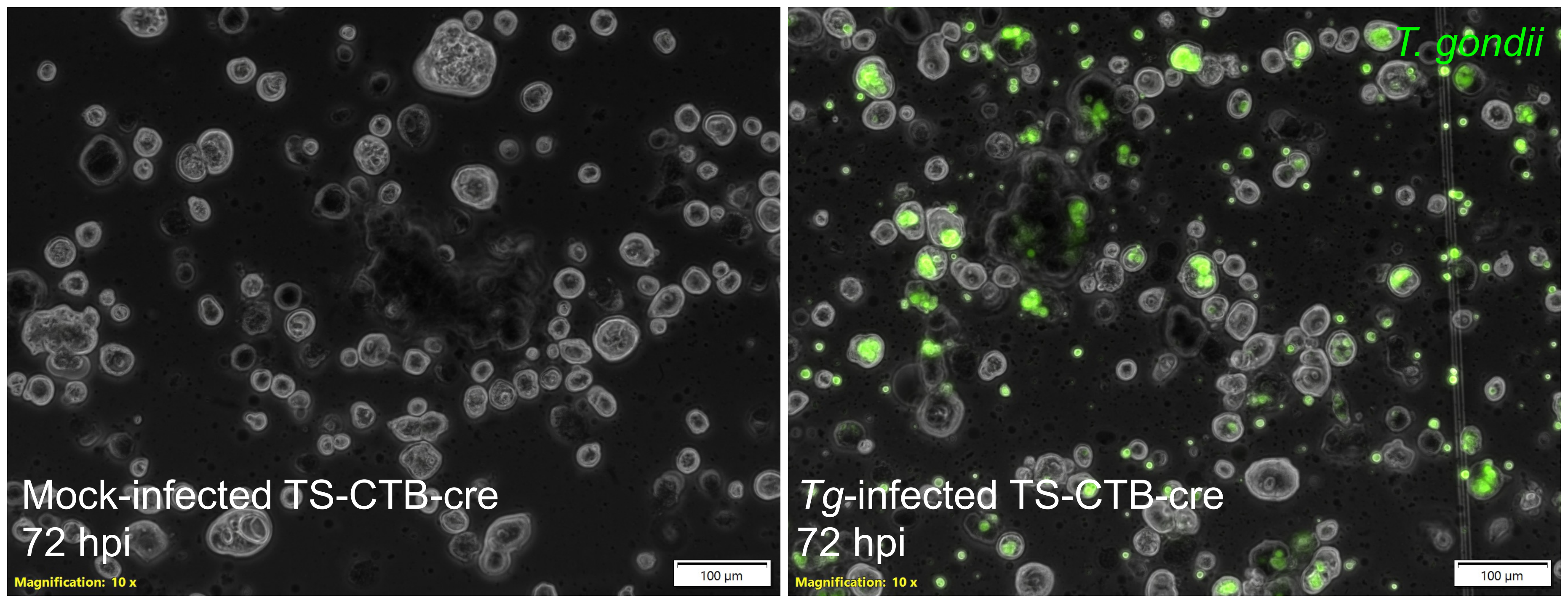


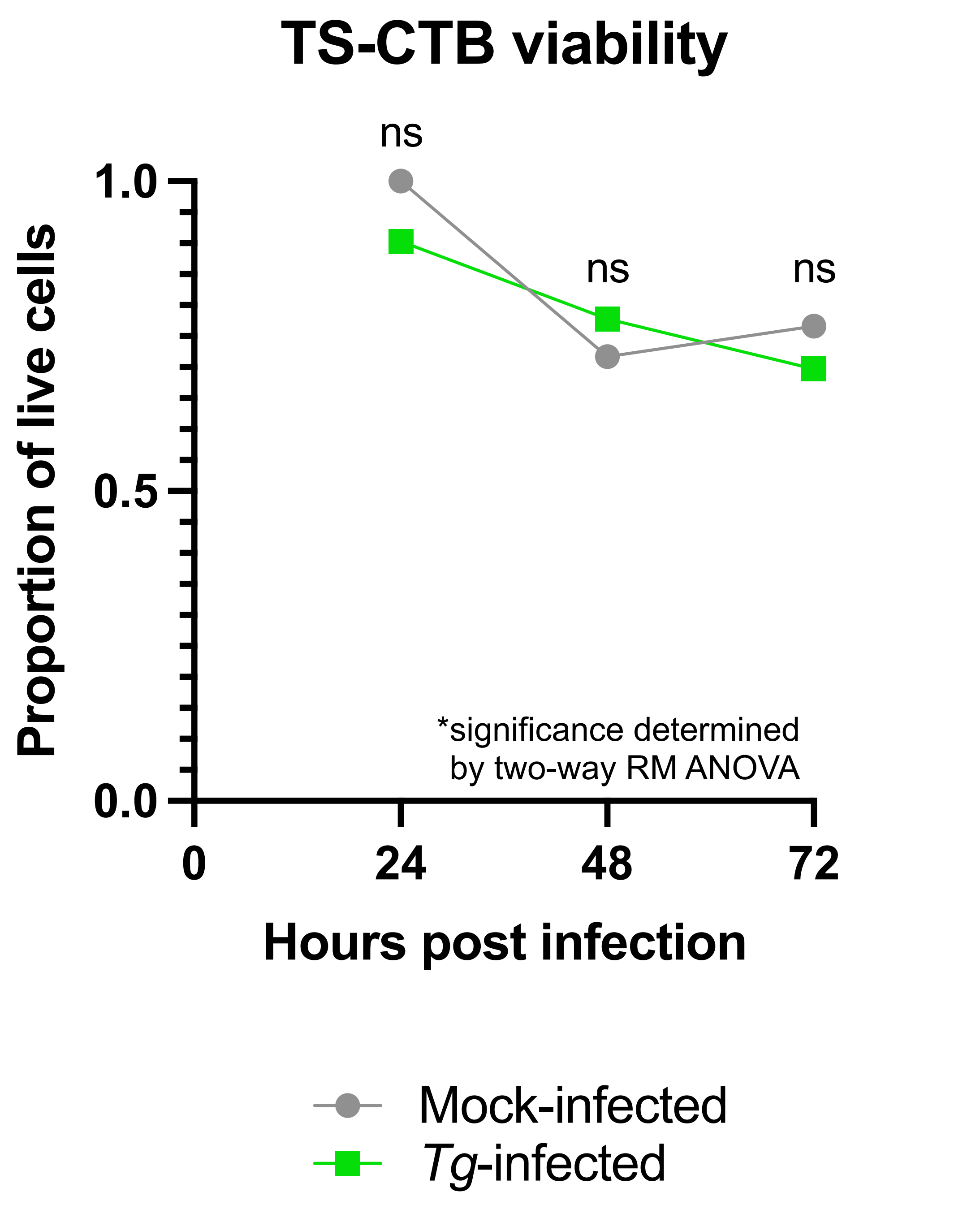

Click here to access/download;Supplemental Figure;figureS2.pdf

### Figure S2

**A**

| | Homo sapiens (REF) | upload_1 (Hierarchy) NEW! | | | | | |
|---|---|---|---|---|---|---|---|
| GO biological process complete | # | # | expected | Fold Enrichment | +/- | raw P value | FDR |
| anatomical structure morphogenesis | 2234 | 30 | 9.23 | 3.25 | + | 2.02E-09 | 6.15E-06 |
| external encapsulating structure organization | 279 | 12 | 1.15 | 10.41 | + | 1.66E-09 | 6.33E-06 |
| extracellular structure organization | 278 | 12 | 1.15 | 10.45 | + | 1.60E-09 | 8.11E-06 |
| animal organ development | 2863 | 35 | 11.82 | 2.96 | + | 6.03E-10 | 9.19E-06 |
| extracellular matrix organization | 277 | 12 | 1.14 | 10.49 | + | 1.53E-09 | 1.17E-05 |
| circulatory system development | 928 | 19 | 3.83 | 4.96 | + | 5.66E-09 | 1.23E-05 |
| tissue development | 1762 | 26 | 7.28 | 3.57 | + | 5.10E-09 | 1.30E-05 |
| regulation of cell population proliferation | 1673 | 25 | 6.91 | 3.62 | + | 8.59E-09 | 1.45E-05 |
| positive regulation of cell population proliferation | 947 | 19 | 3.91 | 4.86 | + | 7.87E-09 | 1.50E-05 |
| regulation of locomotion | 1041 | 19 | 4.30 | 4.42 | + | 3.58E-08 | 4.20E-05 |
| blood vessel development | 530 | 14 | 2.19 | 6.40 | + | 3.38E-08 | 4.30E-05 |
| regulation of cell migration | 939 | 18 | 3.88 | 4.64 | + | 4.18E-08 | 4.56E-05 |
| multicellular organism development | 3957 | 39 | 16.34 | 2.39 | + | 3.29E-08 | 4.56E-05 |
| system development | 3541 | 36 | 14.63 | 2.46 | + | 4.63E-08 | 4.70E-05 |
| developmental process | 5723 | 48 | 23.64 | 2.03 | + | 3.16E-08 | 4.82E-05 |
| vasculature development | 551 | 14 | 2.28 | 6.15 | + | 5.48E-08 | 4.91E-05 |
| anatomical structure development | 5216 | 45 | 21.54 | 2.09 | + | 5.43E-08 | 5.17E-05 |
| response to organic substance | 2465 | 29 | 10.18 | 2.85 | + | 8.16E-08 | 6.91E-05 |
| regulation of cell motility | 999 | 18 | 4.13 | 4.36 | + | 1.06E-07 | 8.53E-05 |
| positive regulation of cellular metabolic process | 3328 | 34 | 13.75 | 2.47 | + | 1.23E-07 | 9.34E-05 |
| cellular developmental process | 3646 | 35 | 15.06 | 2.32 | + | 4.03E-07 | 2.79E-04 |
| cell differentiation | 3643 | 35 | 15.05 | 2.33 | + | 3.97E-07 | 2.88E-04 |
| tissue morphogenesis | 574 | 13 | 2.37 | 5.48 | + | 6.35E-07 | 4.20E-04 |
| collagen catabolic process | 41 | 5 | .17 | 29.53 | + | 7.11E-07 | 4.52E-04 |
| tube development | 904 | 16 | 3.73 | 4.29 | + | 7.69E-07 | 4.69E-04 |
| response to endogenous stimulus | 1410 | 20 | 5.82 | 3.43 | + | 8.66E-07 | 5.08E-04 |
| regulation of multicellular organismal process | 2959 | 30 | 12.22 | 2.45 | + | 1.17E-06 | 6.62E-04 |
| muscle structure development | 516 | 12 | 2.13 | 5.63 | + | 1.36E-06 | 7.38E-04 |
| blood vessel morphogenesis | 431 | 11 | 1.78 | 6.18 | + | 1.59E-06 | 8.34E-04 |
| anatomical structure formation involved in morphogenesis | 969 | 16 | 4.00 | 4.00 | + | 1.91E-06 | 9.68E-04 |

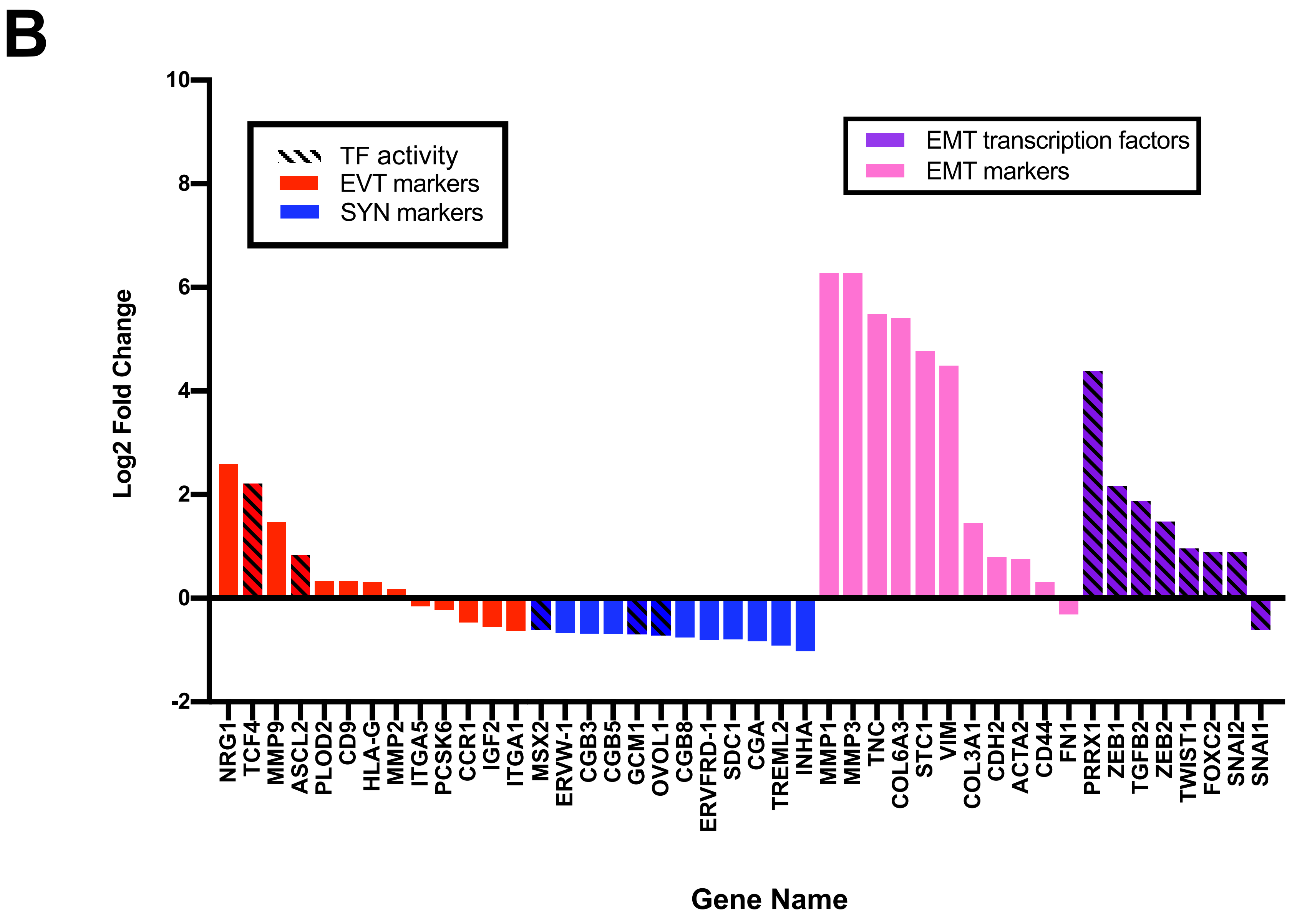


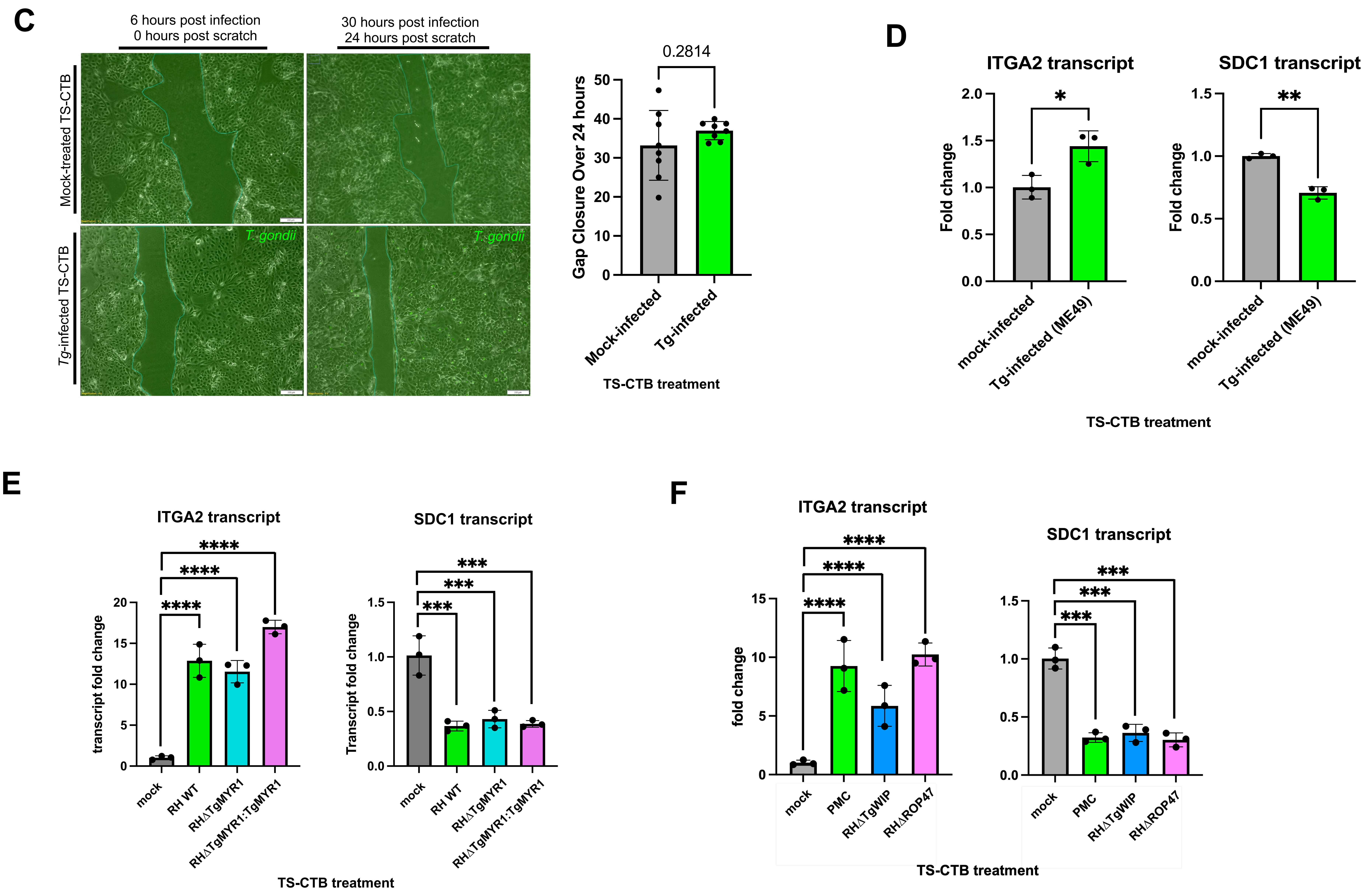

Supplemental Figure 3
Click here to access/download;Supplemental Figure;figureS3.pdf

### Figure S3

A

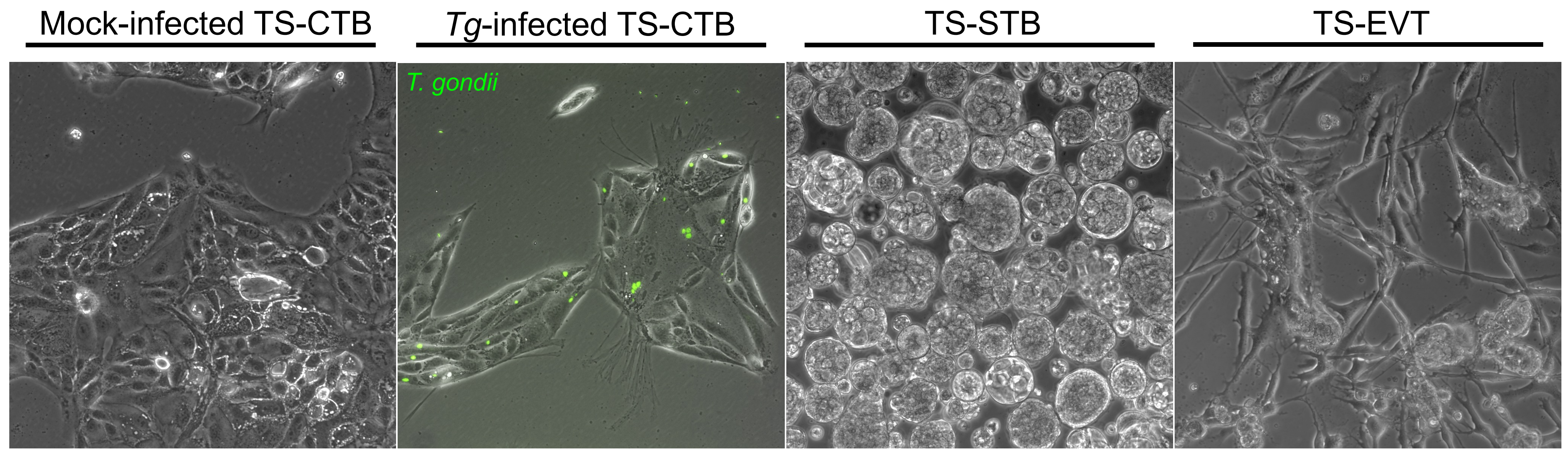


B

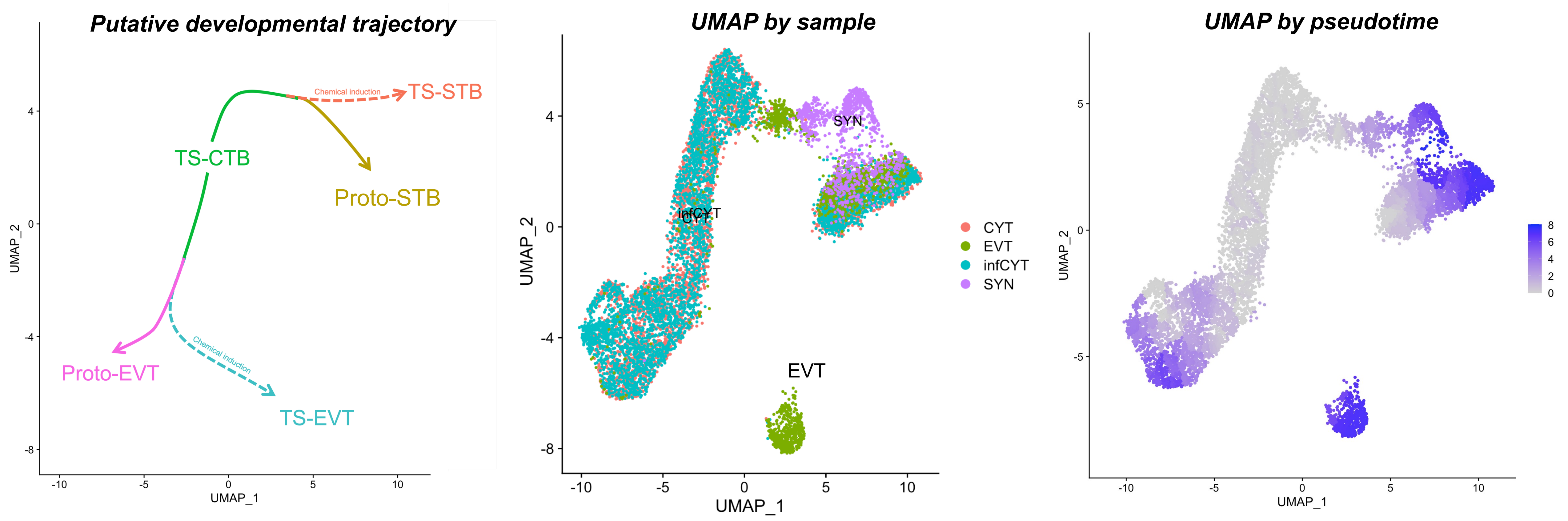


C

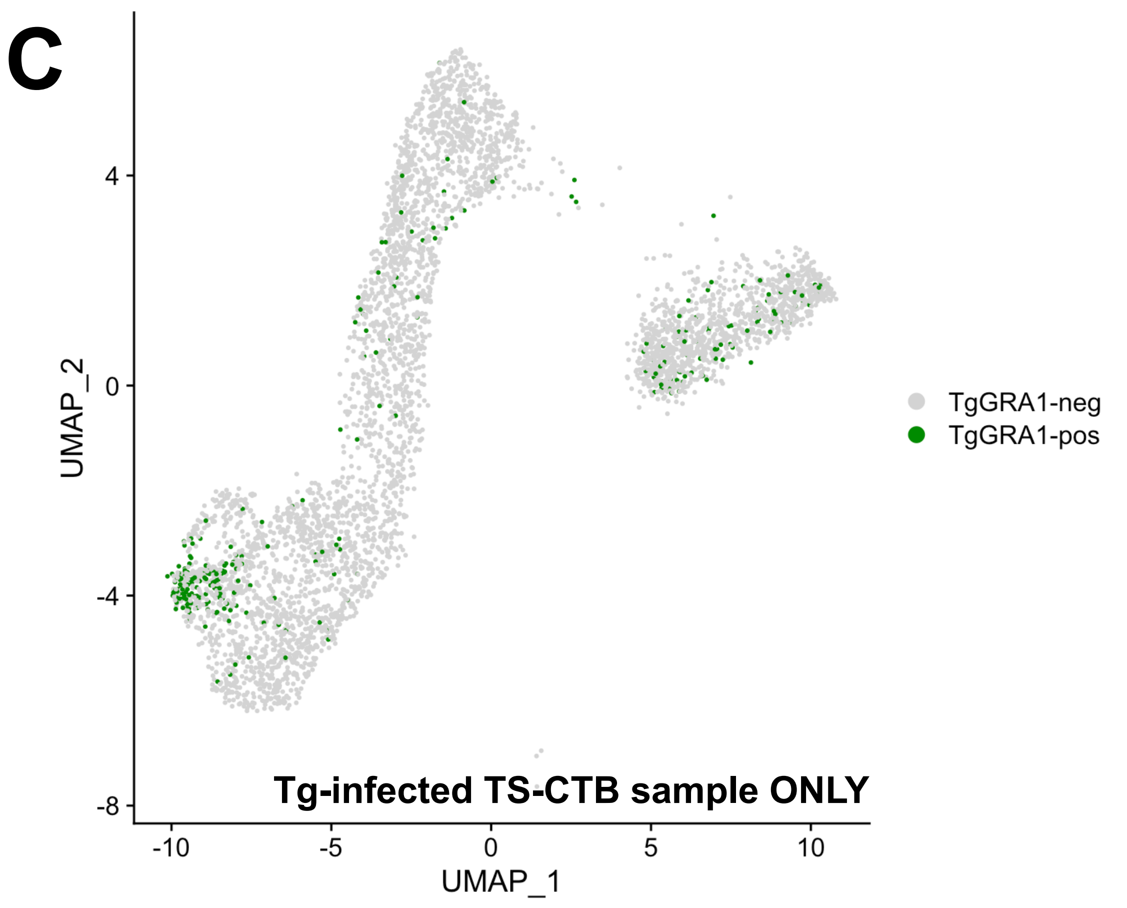


D

Top 10 increased transcripts in TgGRA1(+) cells in TS-EVT predisposed cluster

| Rank | Gene | log2FC | Adj p-val |
|---|---|---|---|
| 1 | BACH2 | 2.12 | 2.63E-12 |
| 2 | DLG2 | 2.05 | 0.00012285 |
| 3 | DDAH1 | 2.02 | 7.69E-15 |
| 4 | SH3RF2 | 1.76 | 2.04E-10 |
| 5 | SPTBN1 | 1.75 | 1.10E-13 |
| 6 | SAMD4A | 1.75 | 8.74E-11 |
| 7 | LMCD1 | 1.73 | 8.71E-09 |
| 8 | ITGA2 | 1.68 | 8.27E-07 |
| 9 | PALLD | 1.64 | 2.83E-11 |
| 10 | BCAT1 | 1.57 | 9.57E-12 |

*ranked by log2FC, filtered p < 0.05

Bottom 10 decreased genes in TgGRA1(+) cells in TS-EVT predisposed cluster

| Rank | Gene | log2FC | Adj p-val |
|---|---|---|---|
| 266 | CGA | -3.68 | 3.60E-36 |
| 265 | ADAMTS20 | -3.32 | 3.74E-31 |
| 264 | CYP19A1 | -3.14 | 4.65E-35 |
| 263 | CGB3 | -2.73 | 2.11E-20 |
| 262 | MYCNUT | -2.53 | 1.05E-23 |
| 261 | NECTIN3 | -2.47 | 7.24E-19 |
| 260 | LINC02506 | -2.35 | 1.32E-19 |
| 259 | ADAM12 | -2.23 | 2.68E-13 |
| 258 | CBLB | -2.22 | 3.62E-26 |
| 257 | CPM | -2.19 | 1.04E-15 |

*ranked by log2FC, filtered p < 0.05

E

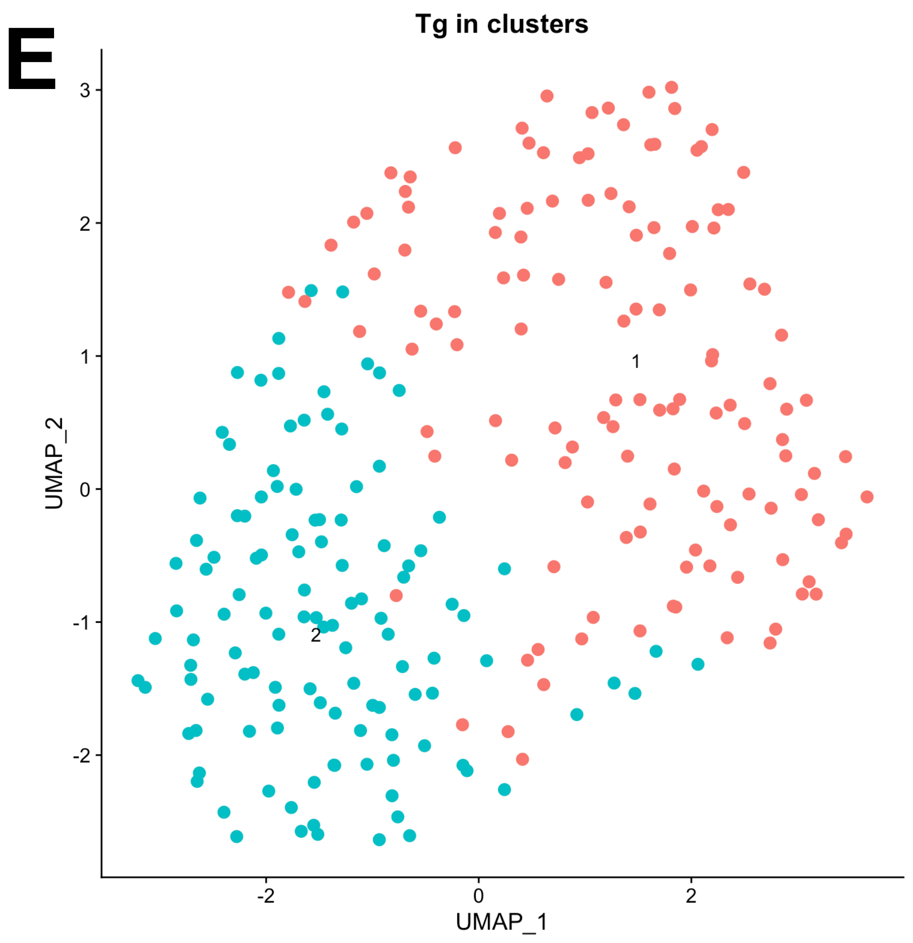


F

All significantly differentially expressed transcripts between clusters 1 and 2 (n = 12)

| Rank | Gene | ToxoDB annotation | log2FC | Adj p-val |
|---|---|---|---|---|
| 1 | TGME49_300100 | rhoptry neck protein RON2 | 2.64 | 5.4E-05 |
| 2 | TGME49_297960 | rhoptry neck protein RON6 | 1.99 | 3.7E-03 |
| 3 | TGME49_293170 | CH domain-containing protein | 1.92 | 1.1E-02 |
| 4 | TGME49_311470 | rhoptry neck protein RON5 | 1.92 | 7.9E-05 |
| 5 | TGME49_305160 | histone H2Ba | 1.86 | 1.2E-05 |
| 6 | TGME49_300220 | hypothetical protein | 1.02 | 2.1E-02 |
| 7 | TGME49_315210 | rhoptry protein, putative | 1.01 | 2.0E-03 |
| 8 | TGME49_306060 | rhoptry neck protein RON8 | 0.96 | 2.7E-03 |
| 9 | TGME49_311230 | basal complex component BCC7 | 0.92 | 4.2E-02 |
| 10 | TGME49_239260 | histone H4 | 0.92 | 3.6E-02 |
| 11 | TGME49_229010 | rhoptry neck protein RON4 | 0.86 | 1.6E-02 |
| 12 | TGME49_206430 | formin FRM1 | 0.70 | 1.8E-02 |

G

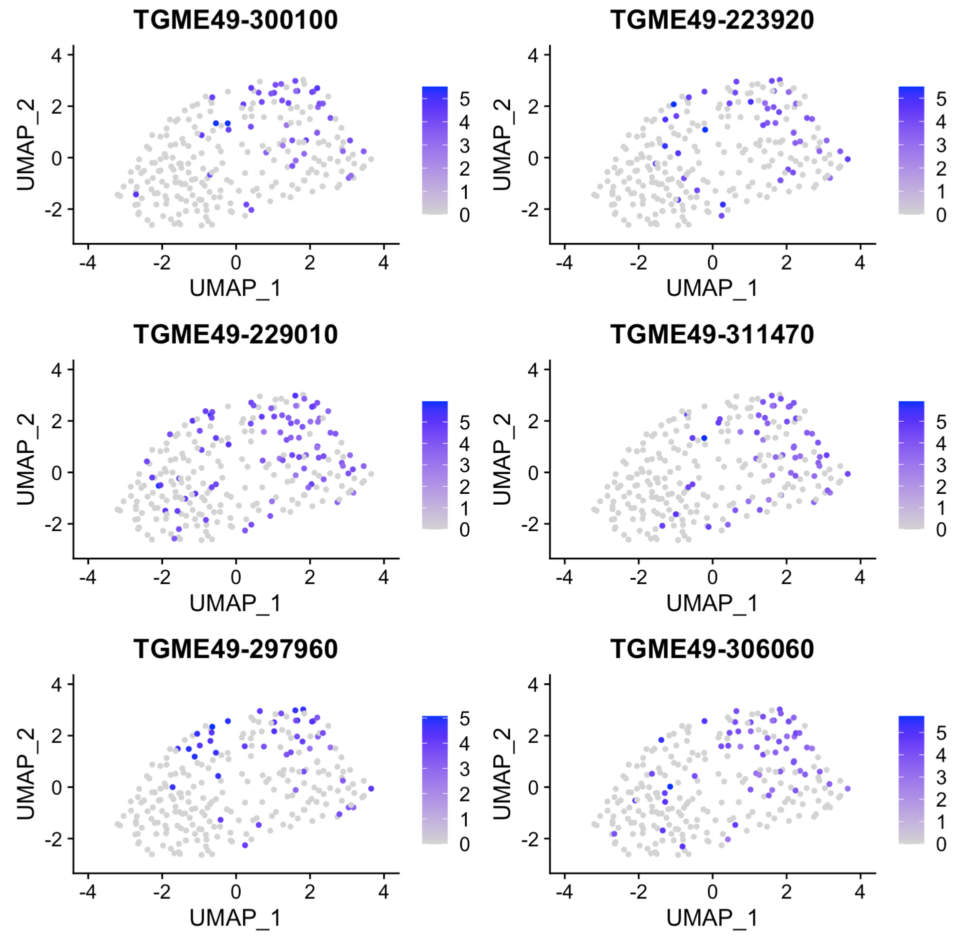

Click here to access/download;Supplemental Figure;figureS4.pdf

### Figure S4

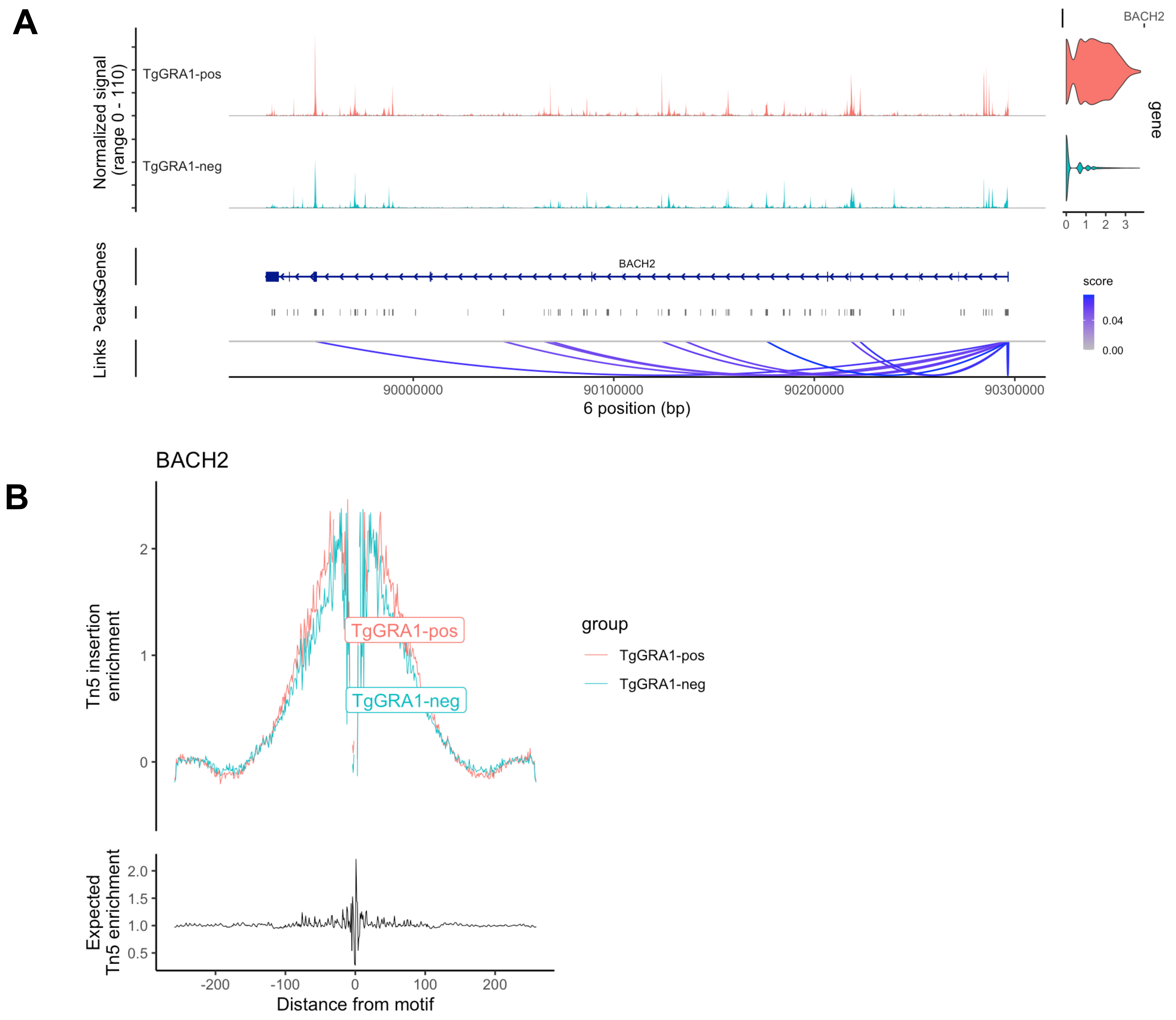

Click here to access/download;Supplemental Figure;figureS5.pdf

### Figure S5

| Gene sets enriched in Tg-infected CV versus mock-infected CV | | | | | | |
|---|---|---|---|---|---|---|
| Rank | Gene Set (GS) | GS size | Normalized enrichment score | Nominal p-value | FDR q-value | FWER p-value |
| 1 | Hallmark_KRAS_Signaling_Up | 198 | 1.57 | 0.000 | 0.268 | 0.219 |
| 2 | Hallmark_Apoptosis | 160 | 1.46 | 0.105 | 6.26 | 0.415 |
| 3 | Hallmark_TNFa_Signaling_via_NFKb | 198 | 1.44 | 0.000 | 0.590 | 0.415 |
| 4 | Hallmark_MYC_Targets_V2 | 57 | 1.44 | 0.000 | 0.497 | 0.415 |
| 5 | Hallmark_IL2_STAT5_Signaling | 197 | 1.42 | 0.208 | 0.475 | 0.466 |
| 6 | Hallmark_Hypoxia | 197 | 1.41 | 0.105 | 0.446 | 0.466 |
| 7 | Hallmark_UV_Response_Up | 156 | 1.40 | 0.097 | 0.410 | 0.466 |
| 8 | Hallmark_Inflammatory_Response | 200 | 1.37 | 0.000 | 0.445 | 0.507 |
| 9 | Hallmark_Bile_Acid_Metabolism | 112 | 1.35 | 0.000 | 0.423 | 0.507 |
| 10 | Hallmark_Reactive_Oxygen_Species_Pathway | 49 | 1.34 | 0.110 | 0.456 | 0.507 |
| 11 | Hallmark_IL6_JAK_STAT3_Signaling | 87 | 1.32 | 0.000 | 0.483 | 0.558 |
| 12 | Hallmark_Allograft_Rejection | 196 | 1.30 | 0.000 | 0.481 | 0.558 |
| 13 | Hallmark_WNT_Beta_Catenin_Signaling | 42 | 1.30 | 0.323 | 0.460 | 0.558 |
| 14 | Hallmark_MTORC1_Signaling | 195 | 1.29 | 0.308 | 0.455 | 0.594 |
| 15 | Hallmark_Xenobiotic_Metabolism | 197 | 1.28 | 0.000 | 0.446 | 0.653 |
| 16 | Hallmark_Oxidative_Phosphorylation | 185 | 1.24 | 0.203 | 0.466 | 0.653 |
| 17 | Hallmark_MYC_Targets_V1 | 195 | 1.24 | 0.203 | 0.474 | 0.704 |
| 18 | Hallmark_Unfolded_Protein_Response | 108 | 1.19 | 0.308 | 0.522 | 0.704 |
| 19 | Hallmark_TGFbeta_Signaling | 54 | 1.19 | 0.308 | 0.503 | 0.764 |
| 20 | Hallmark_Complement | 200 | 1.17 | 0.403 | 0.506 | 0.764 |

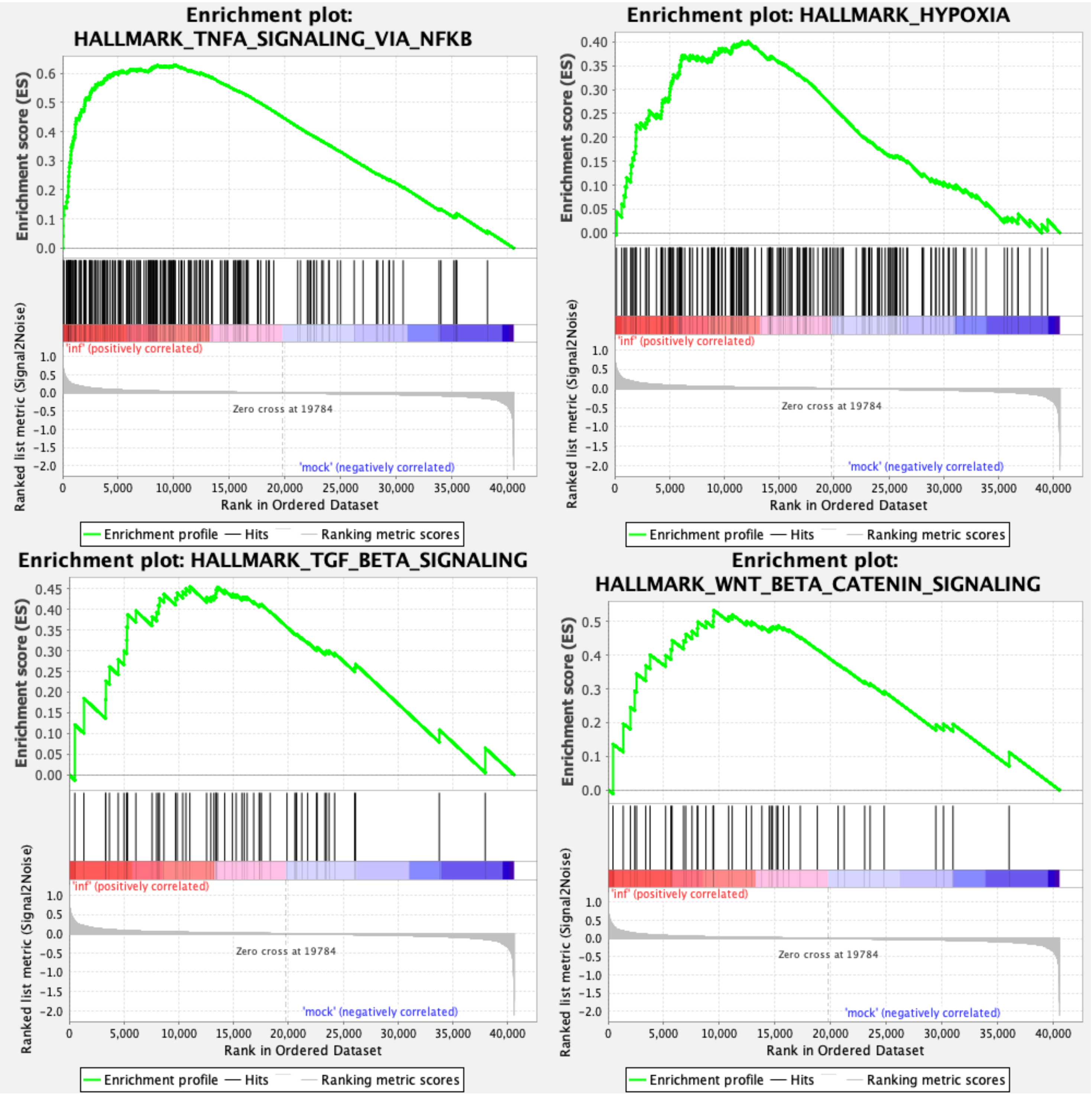

Supplemental Figure 6
Click here to access/download;Supplemental Figure;figureS6.pdf

### Figure S6

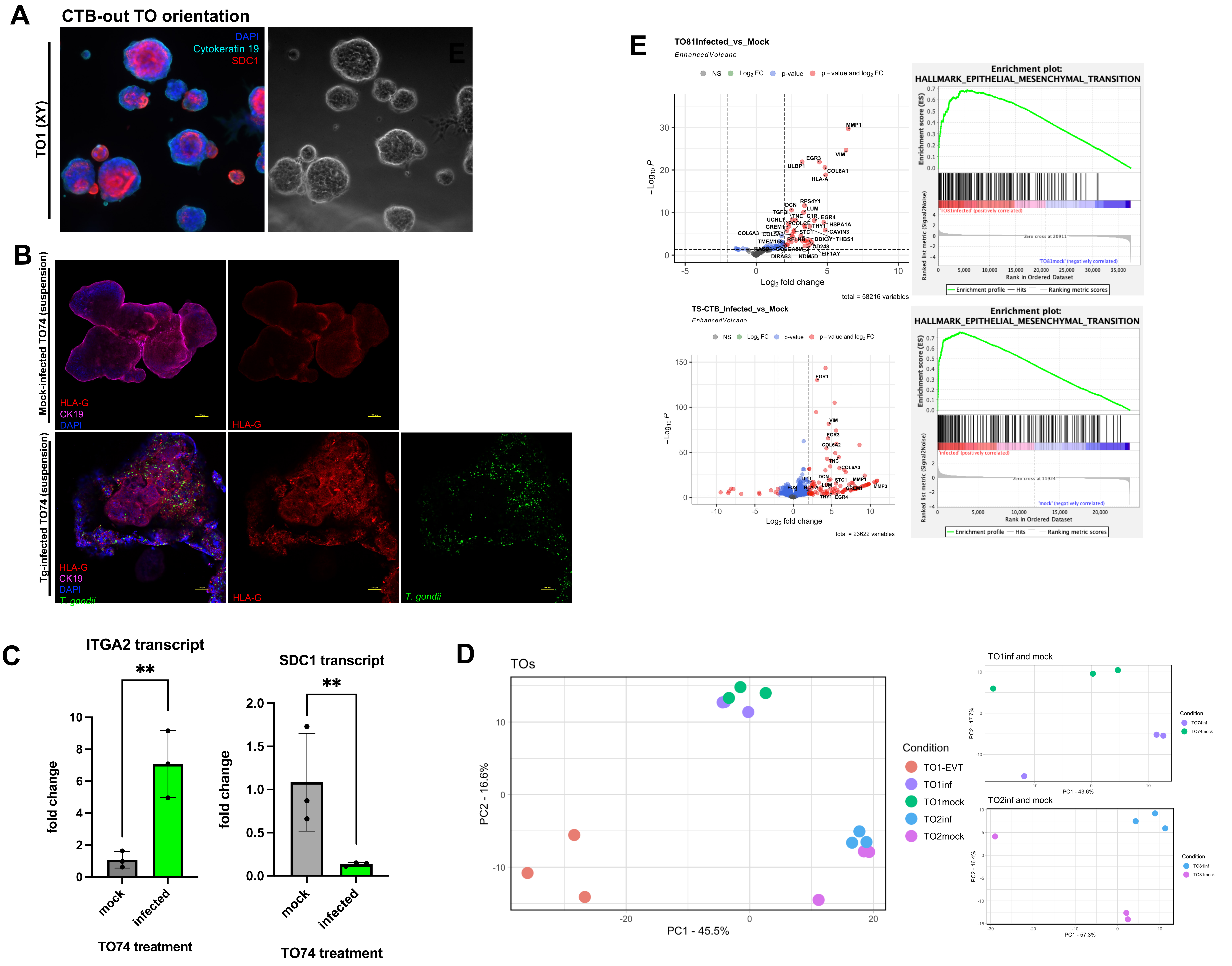

Supplemental Figure 7

Click here to access/download;Supplemental Figure;figureS7.pdf

### Figure S7

A

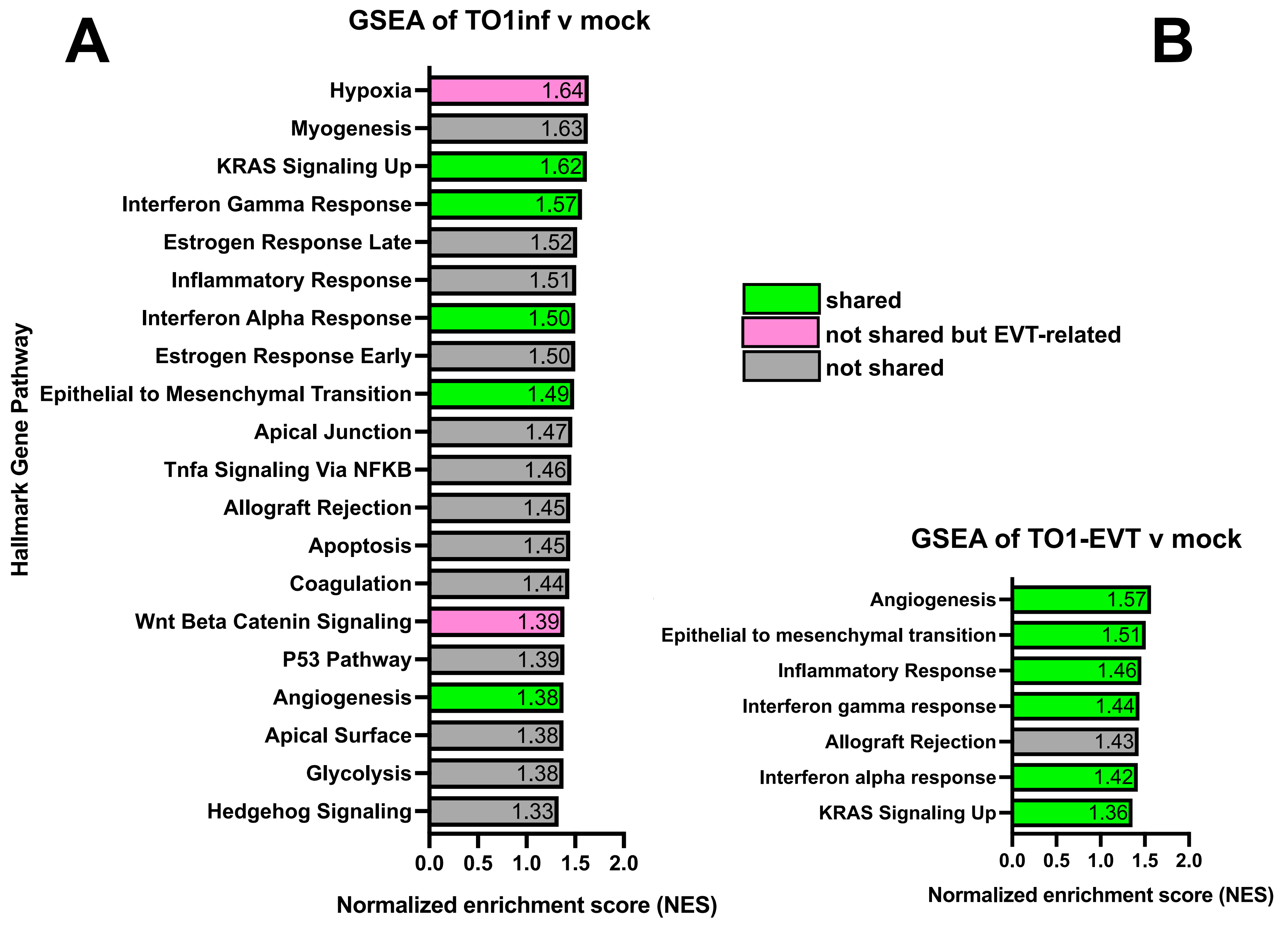


B

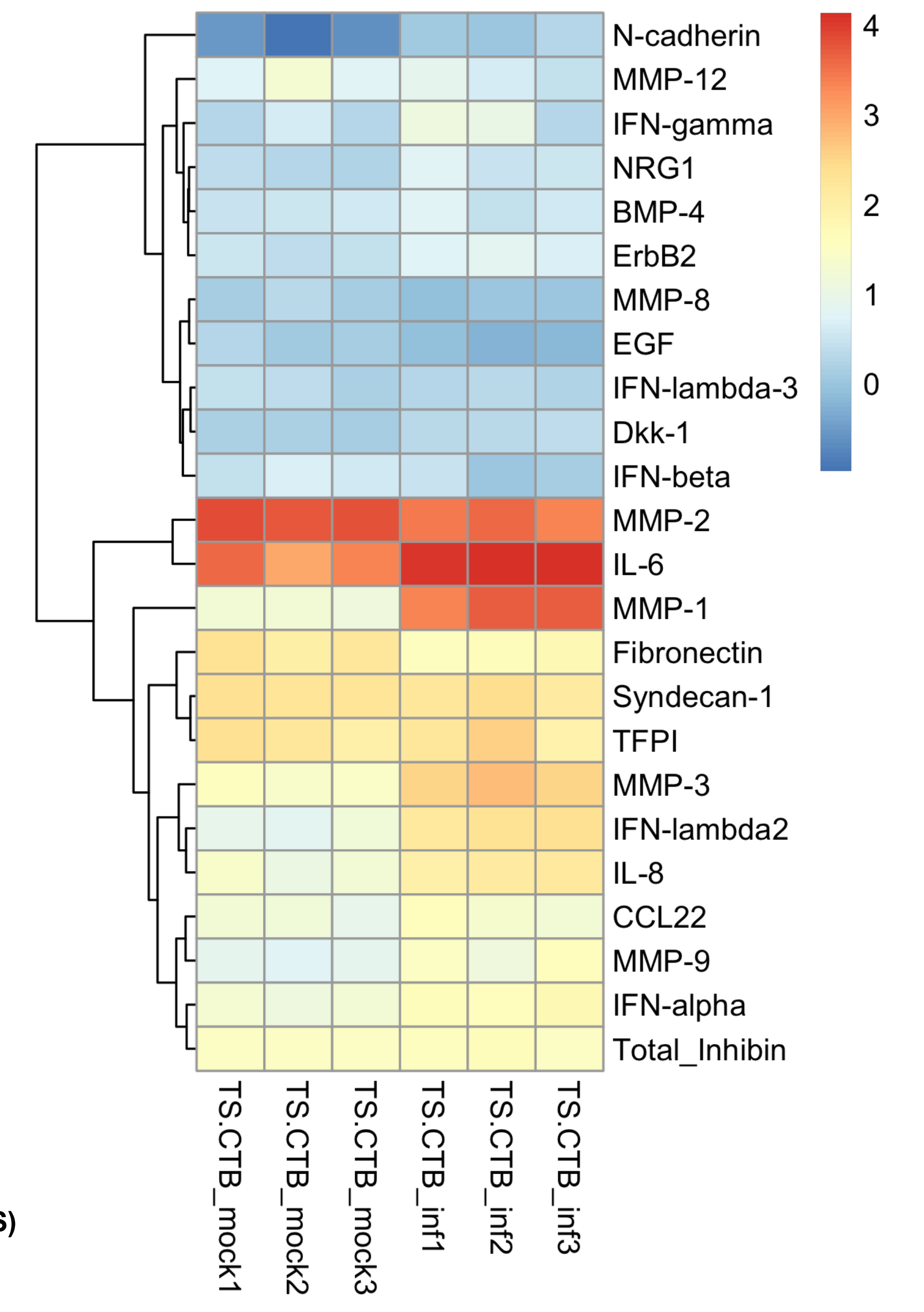


C

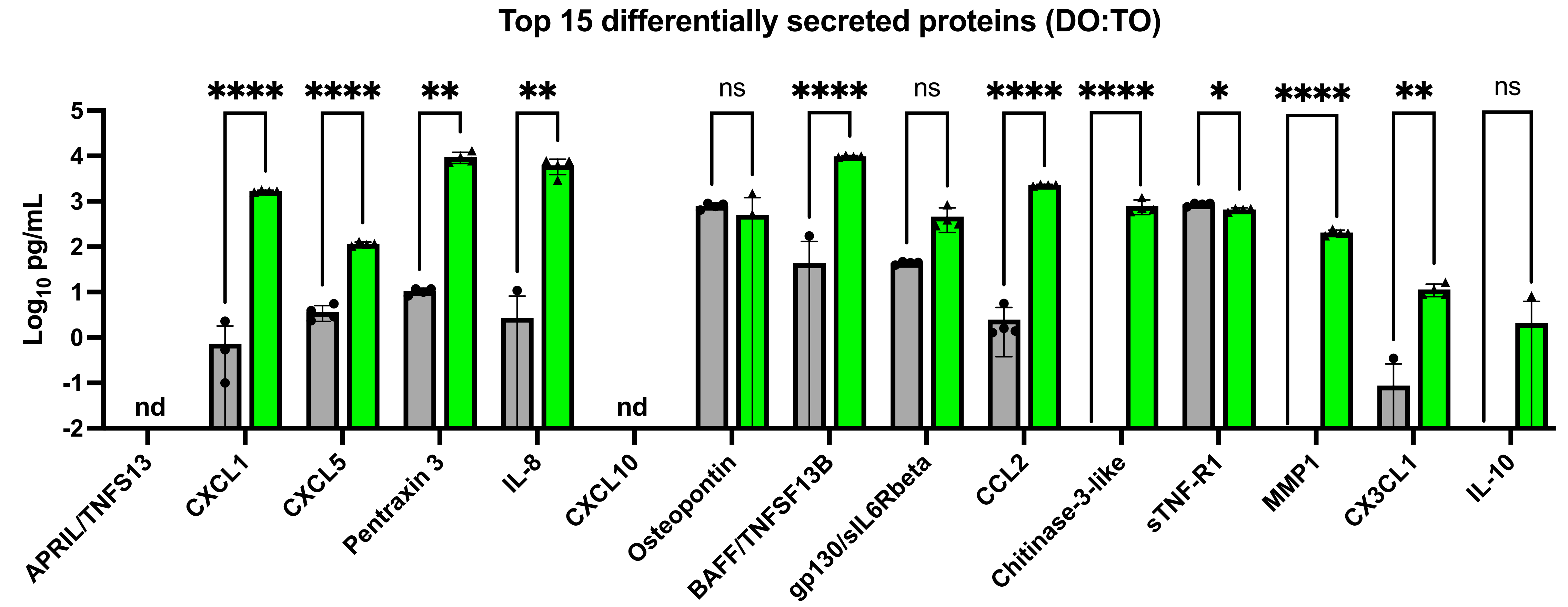


D

| Dataset: | TS-CTB (sn) | | TS-CTB (bulk) | | TO74 (bulk) | | PHT (bulk) | | CV (bulk) | | All |
|---|---|---|---|---|---|---|---|---|---|---|---|
| Gene | Log2 fold change | P-value | Log2 fold change | P-value | Log2 fold change | P-value | Log2 fold change | P-value | Log2 fold change | P-value | (+) fold change with sig p-value |
| *EGR1* | +1.17 | **** | +3.06 | **** | +3.40 | **** | +4.01 | **** | +2.15 | **** | *5/5* |
| *EGR3* | +2.99 | **** | +4.40 | **** | +5.08 | **** | +9.60 | **** | +3.96 | **** | *5/5* |
| *TIMP1* | +0.48 | **** | +0.66 | **** | +0.91 | * | +0.69 | **** | +1.36 | **** | *5/5* |
| *IL6* | +0.75 | **** | +1.45 | **** | - | *ns* | +0.91 | *** | +1.92 | **** | *4/5* |
| *MMP1* | - | *ns* | +7.74 | **** | +6.29 | **** | - | *ns* | +2.95 | **** | *3/5* |
| *MMP3* | - | *ns* | +7.44 | **** | +6.69 | **** | - | *ns* | +4.25 | **** | *3/5* |
| *GREM1* | - | *ns* | +5.92 | **** | +2.63 | **** | -0.93 | **** | +1.82 | **** | *3/5* |
| *DKK1* | +3.99 | **** | - | *ns* | +2.64 | ** | - | *ns* | +2.21 | **** | *3/5* |
| *BACH2* | +2.33 | **** | +0.29 | * | - | *ns* | +1.35 | **** | - | *ns* | *3/5* |
| *INHBA* | +3.16 | **** | +2.06 | **** | - | *ns* | +1.28 | **** | - | *ns* | *3/5* |
| *LIF* | +0.98 | **** | - | *ns* | - | *ns* | +1.98 | **** | +2.69 | **** | *3/5* |
| *TGFB2* | +2.22 | **** | +1.87 | **** | - | *ns* | +0.91 | **** | - | *ns* | *3/5* |
| *NRG1* | +3.53 | **** | +2.59 | **** | - | *ns* | - | *ns* | - | *ns* | *2/5* |
| *CXCL1* | - | *ns* | +0.50 | * | - | *ns* | - | *ns* | +1.12 | * | *2/5* |
| *CXCL8* | - | *ns* | +6.21 | **** | - | *ns* | -1.35 | **** | +2.10 | **** | *2/5* |
| *MMP9* | +2.39 | **** | +1.13 | **** | - | *ns* | -1.22 | **** | -0.60 | * | *2/5* |
| *MMP10* | - | *ns* | +7.93 | **** | - | *ns* | - | *ns* | +3.90 | **** | *2/5* |
| *VEGFA* | - | *ns* | - | *ns* | - | *ns* | +0.75 | **** | +0.80 | **** | *2/5* |
| *STAT3* | +0.47 | **** | -0.27 | ** | - | *ns* | +0.51 | **** | - | *ns* | *2/5* |

E

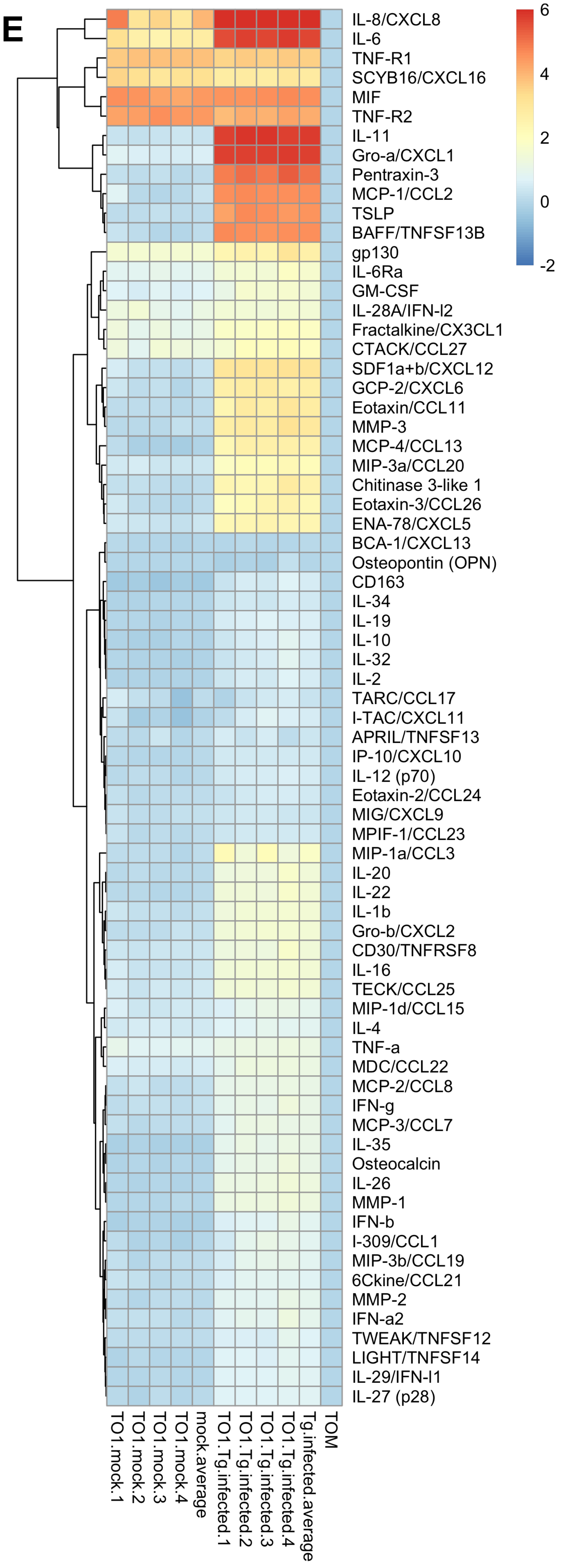
